# Pairing-dependent plasticity in a dissected fly brain is input-specific and requires synaptic CaMKII enrichment and nighttime sleep

**DOI:** 10.1101/2022.02.08.479633

**Authors:** Mohamed Adel, Nannan Chen, Yunpeng Zhang, Martha L. Reed, Christina Quasney, Leslie C. Griffith

**Affiliations:** Department of Biology and Volen National Center for Complex Systems, Brandeis University, Waltham, MA 02454-9110, USA

## Abstract

In *Drosophila*, *in vivo* functional imaging studies revealed that associative memory formation is coupled to a cascade of neural plasticity events in distinct compartments of the mushroom body (MB). In-depth investigation of the circuit dynamics, however, will require an *ex vivo* model that faithfully mirrors these events to allow direct manipulations of circuit elements that are inaccessible in the intact fly. The current *ex vivo* models have been able to reproduce the fundamental plasticity of aversive short-term memory, a potentiation of the MB intrinsic neurons (Kenyon cells; KCs) responses after artificial learning *ex vivo*. However, this potentiation showed different localization and encoding properties from those reported *in vivo* and failed to generate the previously reported suppression plasticity in the mushroom body output neurons (MBONs). Here, we develop an *ex vivo* model using the female *Drosophila* brain that recapitulates behaviorally evoked plasticity in the KCs and MBONs. We demonstrate that this plasticity accurately localizes to the MB α’3 compartment and is encoded by a coincidence between KCs activation and dopaminergic input. The formed plasticity is input-specific, requiring pairing of the conditioned stimulus (CS) and unconditioned stimulus (US) pathways; hence we name it pairing-dependent plasticity (PDP). PDP formation requires an intact *CaMKII* gene and is blocked by previous-night sleep deprivation but is rescued by rebound sleep. In conclusion, we show that our *ex vivo* preparation recapitulates behavioral and imaging results from intact animals and can provide new insights into mechanisms of memory formation at the level of molecules, circuits, and brain state.

**Significance Statement:** The mammalian *ex vivo* LTP model enabled in-depth investigation of the hippocampal memory circuit. We develop a parallel model to study the *Drosophila* mushroom body (MB) memory circuit. Pairing activation of the conditioned stimulus and unconditioned stimulus pathways in dissected brains induces a potentiation pairing-dependent plasticity (PDP) in the axons of α’β’ Kenyon cells and a suppression PDP in the dendrites of their postsynaptic MB output neurons, localized in the MB α’3 compartment. This PDP is input-specific and requires the 3’ untranslated region of *CaMKII*. Interestingly, *ex vivo* PDP carries information about the animal’s experience before dissection; brains from sleep deprived animals fail to form PDP while those from animals who recovered 2 hours of their lost sleep form PDP.

## Introduction

A neutral experience (conditioned stimulus or CS) can be remembered as positive or negative if closely followed by rewarding or punishing reinforcement (unconditioned stimulus or US). The ability to form this type of “associative” memory is phylogenetically conserved; *Drosophila* form robust associative memories (Tully and Quinn, 1985), most of which are encoded and stored in the mushroom body (MB) (de Belle and Heisenberg, 1994). The MB is a higher brain structure made of 15 distinct compartments. Each compartment is built on a scaffold of axons of one of the three main types of Kenyon cells (KCs; αβ, α’β’, and γ). The KCs connect to mushroom body output neurons (MBONs) which project out of the MB to bias behavior (Aso et al., 2014a; Aso et al., 2014b; Li et al., 2020). The KC→MBON synapses are modulated by dopaminergic neurons.

During aversive olfactory associative learning, an odor (the CS) activates a sparse group of KCs, such that this odor identity is represented across all MB compartments (Turner et al., 2008; Lin et al., 2014). Simultaneously, dopaminergic neurons from the protocerebral posterior lateral (PPL1) cluster are activated by the US, encoding negative prediction errors in MB compartments (Riemensperger et al., 2005; Schroll et al., 2006; Claridge-Chang et al., 2009; Mao and Davis, 2009; Aso et al., 2012). When KCs activation and the dopaminergic signal coincide within a compartment, the KC MBON synapses in that compartment are depressed, biasing the circuit output to aversion (Sejourne et al., 2011; Cohn et al., 2015; Hige et al., 2015; Owald et al., 2015; Owald and Waddell, 2015).

Many studies have investigated the properties of this circuitry using *in vivo* calcium imaging in intact animals (for review, see Adel and Griffith, 2021). In contrast, explanted brains have been used mostly for establishing connectivity between neurons or interrogating a specific biochemical pathway; only a few studies have attempted to understand memory circuit logic *ex vivo* (Wang et al., 2008; Ueno et al., 2013; Suzuki-Sawano et al., 2017; Ueno et al., 2017). In the best-developed paradigm, Ueno et al., 2013 observed a potentiation of KCs responses in the tips of the MB vertical lobes which they termed “long-term enhancement” (LTE). This laid the groundwork for developing *ex vivo* models of this circuit, but there were major differences between LTE and associative memory observed in intact animals. The most significant were that the plasticity was not specific to the α’β’ KCs and that dopamine release by the US was not observed; it was only seen after CS+US coincidence (Ueno et al., 2013; Ueno et al., 2017).

In this study, we establish an *ex vivo* paradigm that resolves these discrepancies and exhibits the cardinal features of associative learning. We show that pairing odor and punishment pathway activation in dissected brains results in a localized potentiation of the α’β’ KCs and suppression of their postsynaptic MBONs in the α’3 compartment. Because both KC potentiation and MBON suppression are strictly dependent on temporal coincidence of the CS and US, we term this paradigm “pairing-dependent plasticity” (PDP). We show that like the CS-specificity of associative memories, PDP is specific to the subset of odor-representing projection neurons activated during the artificial training. We also provide evidence that dopamine is released by activation of the US pathway and does not require CS+US coincidence.

This *ex vivo* paradigm can be used for obtaining new mechanistic insight into memory formation at the molecular and circuit levels. We present data indicating that the 3’UTR of the *CaMKII* gene is critical for STM formation and that the primacy of α’ compartment plasticity in learning is due to differences in input/response relationships between α and α’. Finally, we demonstrate that the ability of the *ex vivo* brain to be plastic can be influenced by prior *in vivo* experience, as we report that brains of sleep-deprived flies fail to form PDP, but as little as 2 hours of recovery sleep rescues this learning impairment.

## Materials and Methods

### Fly husbandry

All fly stocks were cultured on standard food at room temperature. Experimental flies were kept at 25°C and 70% relative humidity on a 12 hours light, 12 hours dark period. Fly lines used in this study include *VT030559-GAL4, MB027B split-GAL4* (Aso et al., 2014a), *GH146-GAL4, UAS-GCaMP6f, 20x UAS-GCaMP6f, UAS-jRCaMP1a, LexAop-P2X_2_. TH-lexA* was gifted to us from the Davis lab (Berry et al., 2015) and *UAS-GRAB^DA2m^* was gifted from the Li lab (Sun et al., 2020). *CaMKII^UDel^* flies are described in (Chen et al., 2022).

### Ex vivo imaging and electrical stimulations

Brains from 4-8 days old female flies were dissected in ice cold HL3 medium (NaCl, 70 mM; sucrose, 115 mM; KCl, 5 mM; MgCl2, 20 mM; CaCl2, 1.8 mM; NaHCO3, 10 mM; trehalose, 5 mM; Hepes, 5 mM; osmolarity: 395.4 mOsm; pH 7.3) (Stewart et al., 1994). Brains were then transferred to an imaging chamber containing fresh HL3 saline and immobilized using tungsten pins over the optic lobes. In the case of paired stimulations of the antennal lobe (AL) and the ascending fibers of the ventral nerve cord (AFV), the brain was dissected with the ventral nerve cord attached. The ventral nerve cord was then cut using sharp scissors near its base, leaving one end of the AFV free. Glass suction microelectrodes were used to apply the electrical stimulation to either the AL or the AFV. Because odor information is randomly encoded in the MB (Bhandawat et al., 2007; Ito et al., 2008; Turner et al., 2008; Honegger et al., 2011; Lei et al., 2013), we did not target specific AL glomeruli across different animals. However, the properties of the stimulation and the electrode were kept the same across all experiments unless noted otherwise. Based on the size of the AL electrode tip and the distribution of AL calcium responses to AL stimulation (see Figure 3 B-D), we estimate that 20-25% of the ipsilateral AL projection neurons are activated with our AL stimulation protocol. Brains were always perfused with fresh saline throughout the experiment with a flow rate of ∼2 drops per second.

**Figure 1.**
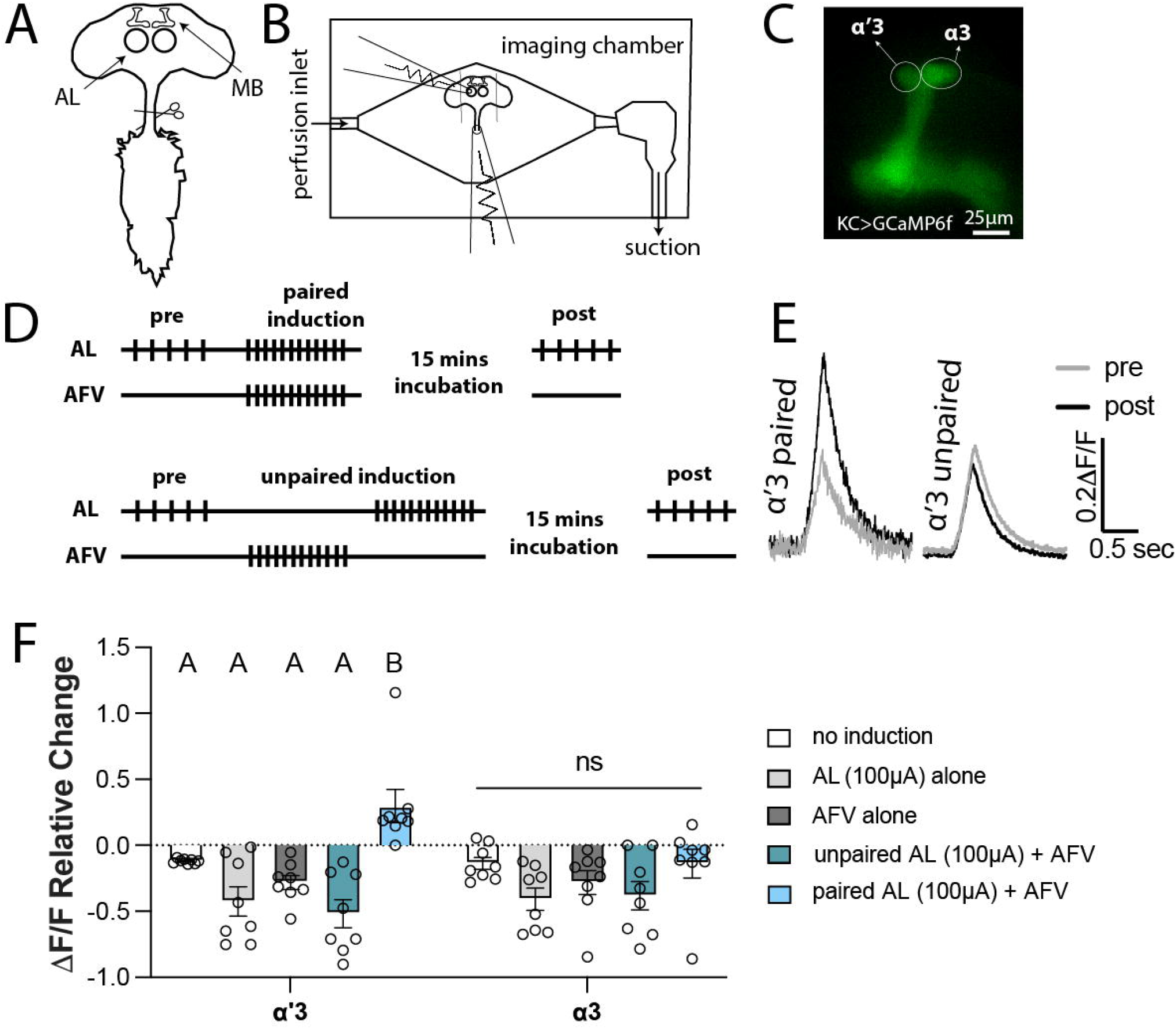
Simultaneous activation of the odor and the electric shock pathways induces an enhancement PDP in the KCs in the MB α’3 compartment. *A,* Schematic of the dissected adult fly’s central nervous system showing the MBs and the ALs in the central brain and the ventral nerve cord. The ventral nerve cord (VNC) is cut at the base of the cervical connective to free the afferent fibers of the VNC (AFV). *B,* Schematic of the imaging chamber showing the placement of the first electrode on the AL, and the second electrode on the free end of the AFV. *C,* Representative image of the MB ipsilateral to the stimulated AL. The calcium indicator GCaMP6f is expressed in the KCs (driven by *VT030559-GAL4*). The circles show the analyzed ROIs surrounding the α’3 and the α3 compartments. Scale bar is 25 μm. *D,* Paired and unpaired induction protocols. Top, in the paired induction, the ipsilateral AL is activated by 5 stimulation trains with 15 seconds intertrain interval followed by 1 minute rest (pre-induction). 12 trains of stimulations are then delivered to both the AL and the AFV simultaneously (induction). The brain is then rested for 15 minutes before being tested by 5 trains of AL stimulations like those applied during the pre-induction (post-induction). Bottom, the unpaired induction: same as paired induction except that the AL and the AFV stimulations are separated by 30 seconds during the induction stage. The stimulation train is 20 pulses at 100 Hz; each pulse is 1 millisecond with 9 milliseconds inter-pulse interval. AL stimulation strength: 100 μAmps. AFV stimulation strength: 0.5-1 mAmps. *E,* Example of the KC calcium response in the α’3 compartment pre (grey) and post (black) paired and unpaired inductions. *F,* Mean relative change of the calcium responses calculated as (*mean post responses – mean pre responses*)/(*mean pre responses*) in the α’3 (left) and the α3 (right) compartments after no induction (white), AL activation alone (light grey), AFV activation alone (dark grey), unpaired AL+AFV induction (green), and paired AL+AFV induction (blue). Mean ±SEM. Two-way ANOVA (α = 0.05; n = 8 in each condition): lobe effects F_(1, 70)_ = 1.096, p= 0.2987; induction effects F_(4,70)_ = 11.51, p<0.0001. Šídák post-hoc tests: In the α’3 compartment: A vs. A, p > 0.05; B vs. A, p ≤ 0.05; t_(70)_ {3.23, p= 0.015; 5.6, p< 0.0001; 4.48, p= 0.0002; 6.319, p<0.0001} for {paired vs. no induction; paired vs. AL alone; paired vs. AFV alone; paired vs. unpaired}, respectively. No statistical significance across conditions in the α3 compartment, p > 0.05; t_(70)_ = {0.0237, p> 0.9999; 2.093, p= 0.2785; 1.105, p = 0.9218; 1.884, p= 0.4094} for {paired vs. no induction; paired vs. AL alone; paired vs. AFV alone; paired vs. unpaired}, respectively.

**Figure 2.**
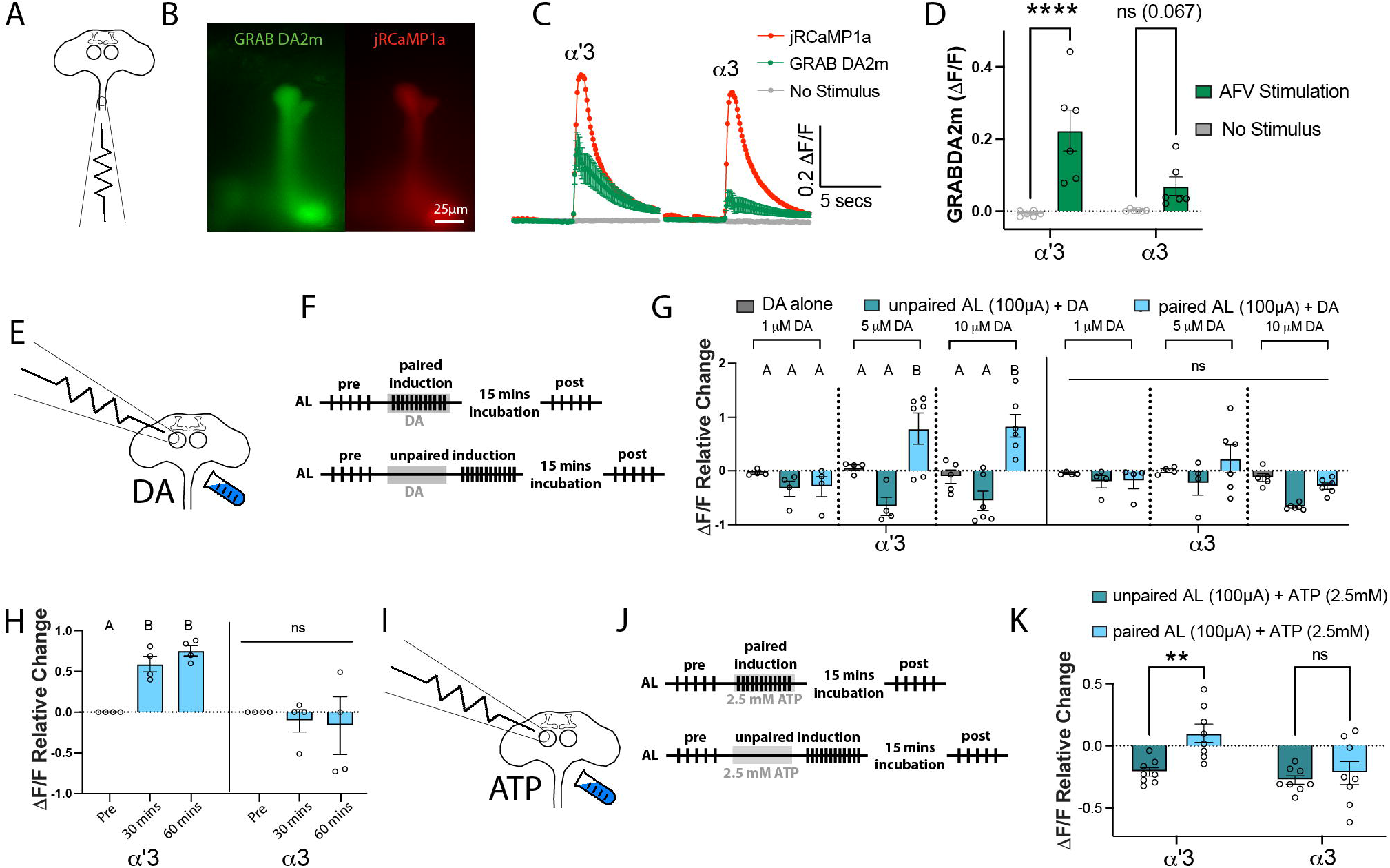
Electric shock pathway activation can be replaced by either dopamine perfusion or activation of dopaminergic neurons. ***A,*** Schematic of the dissected adult central nervous system showing the placement of one electrode to activate the AFV. ***B,*** Representative images showing the simultaneous expression of *UAS-GRAB DA2m* (left; green) and *UAS-jRCaMP1a* (right; red) in the same neurons in the MB (driven by *VT030559-GAL4*). Scale bar is 25 μm. ***C,*** Time course of GRAB DA2m responses in the α’3 and α3 compartments upon AFV stimulation (green) or no stimulus (grey). Traces show mean ΔF/F ±SEM across the 6 flies tested. In each compartment, a representative jRCaMP1a response is shown (red) as a reference for neural activity upon AFV stimulation. ***D,*** Quantification of the GRAB DA2m responses in ***C***. Two-way ANOVA (α = 0.05; n = 6 in each condition): lobe effects F_(1, 20)_ = 3.759, p= 0.0668; stimulation effects F_(1, 20)_ = 29.52, p<0.0001. Šídák post-hoc tests: In the α’3 compartment, ****t_(20)_ = 5.411, p<0.0001; In the α3 compartment, ^ns^t_(20)_ = 2.273, p = 0.0673. ***E,*** Schematic of the AL+DA induction protocol, showing electrode placement above the AL, and perfusion of dopamine. ***F,*** Paired and unpaired induction protocols. Top, in the paired induction, the ipsilateral AL is activated by 5 stimulation trains with 15 seconds intertrain interval followed by resting for 1 minute (pre-induction). 12 trains of stimulations are then applied via the AL electrode, coincident with 60 seconds of dopamine perfusion. The brain is then rested for 15 minutes before being tested by 5 trains of AL stimulations like those applied during the pre-induction (post-induction). Bottom, the unpaired induction: same as paired induction except that the AL stimulation and dopamine perfusion are separated by 30 seconds during the induction stage. The stimulation train is 20 pulses at 100 Hz; each pulse is 1 millisecond with 9 milliseconds inter-pulse interval. AL stimulation strength: 100 μAmps. ***G,*** Mean relative change of the calcium responses in the α’3 (left) and the α3 (right) compartments after DA perfusion alone (grey), unpaired AL+DA induction (green), and paired AL+DA induction (blue). Three different DA concentrations were used: 1 μM (left), 5 μM (middle), and 10 μM (right). Mean ±SEM. Two-way ANOVA: lobe effects F_(1, 68)_ = 2.724, p= 0.1034; induction effects F_(8, 68)_ = 9.584, p<0.0001. Tukey’s post-hoc results are listed in Table 1. ***H,*** Mean relative change of the calcium responses in the α’3 (left) and the α3 (right) compartments pre induction and 30 or 60 minutes after paired AL+ 10 μM DA induction (blue). Mean ±SEM. Two-way ANOVA with repeated measures (α = 0.05; n = 4 in each condition): lobe effects F_(1, 6)_ = 12.67, p= 0.0119; time effects F_(1.341, 8.045)_ = 2.249, p=0.1719; lobe x time interaction effects F_(2,12)_ = 5.206, p=0.0236. Šídák post-hoc tests: in the α’3 compartment: A vs. A, p > 0.05; B vs. A, p ≤ 0.05; t_(3)_ = {6.216, p= 0.025; 11.64, p= 0.0041; 1.03, p= 0.7604} for {pre vs. 30 mins; pre vs. 60 mins; 30 mins vs. 60 mins}, respectively. No statistical significance across conditions in the α3 compartment, p> 0.05; t_(3)_ = {0.7799, p= 0.8692; 0.4625, p= 0.9657; 0.1993, p= 0.9969} for {pre vs. 30 mins; pre vs. 60 mins; 30 mins vs. 60 mins}, respectively. ***I,*** Schematic of the AL+PPL1 induction protocol, showing electrode placement above the AL, and perfusion of 2.5 mM ATP to activate the P2X_2_ channels expressed in the PPL1 dopaminergic neurons (driven by *TH-LexA*). ***J,*** Paired and unpaired induction protocols. Top, in the paired induction, the ipsilateral AL is activated by 5 stimulation trains with 15 seconds intertrain interval followed by resting for 1 minute (pre-induction). We then apply 12 trains of stimulations to the AL electrode, coincident with 60 seconds of 2.5 mM ATP perfusion. The brain is then rested for 15 minutes before being tested by 5 trains of AL stimulations like those applied during the pre-induction (post-induction). Bottom, the unpaired induction: same as paired induction except that the AL stimulation and ATP perfusion are separated by 30 seconds during the induction stage. The stimulation train is 20 pulses at 100 Hz; each pulse is 1 millisecond with 9 milliseconds inter-pulse interval. AL stimulation strength: 100 μAmps. ***K,*** Mean relative change of the calcium responses in the α’3 (left) and the α3 (right) compartments after unpaired AL+ATP induction (green), and paired AL+ATP induction (blue). Mean ±SEM. Two-way ANOVA (α = 0.05; n = 8 in each condition): lobe effects F_(1, 28)_ = 9.053, p= 0.0055; induction effects F_(1, 28)_ = 8.284, p=0.0076. Šídák post-hoc tests: In the α’3 compartment, **t_(28)_ = 3.446, p<0.0036; In the α3 compartment, ^ns^t_(28)_ = 0.624, p = 0.7862.

**Figure 3.**
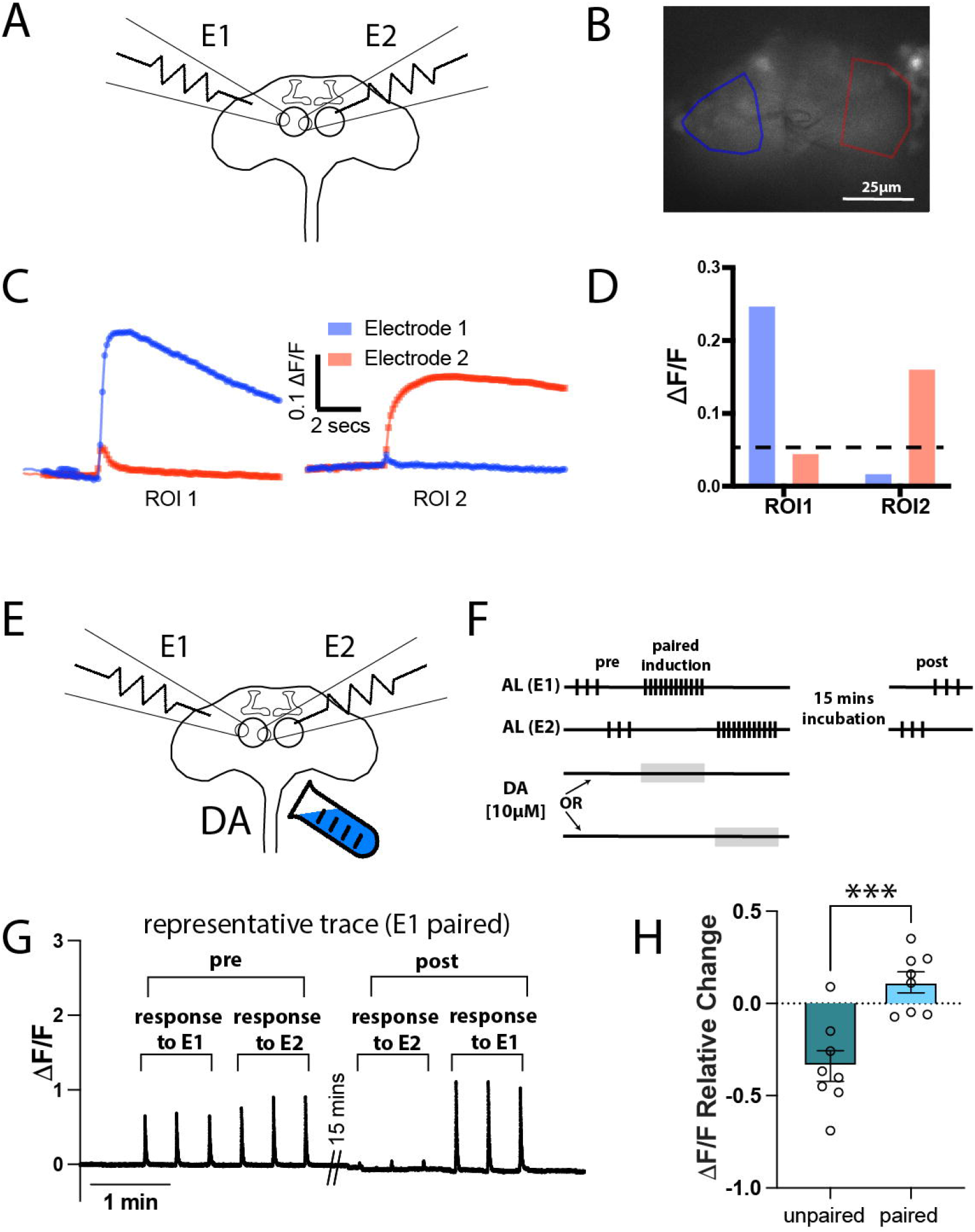
PDP is CS-pathway input-specific. ***A,*** Schematic of the dissected adult brain showing the placement of two electrodes (E1 and E2) onto the surface of the same AL. ***B,*** Basal calcium signals in the AL projection neurons (genotype: *w-;GH146-GAL4/+;UAS-GCaMP6f/+*) showing two ROIs highlighted in blue (ROI near E1) and red (ROI near E2). ***C,*** calcium responses in each ROI to stimulation from E1 (blue trace) or E2 (red trace). ***D,*** Quantification of ***C***. Dashed line is plotted at ΔF/F = 0.05**. *F,*** Induction protocol. The ipsilateral AL is activated by 3 stimulation trains with 15 seconds intertrain interval followed by resting for 1 minute (pre-induction). 12 trains of stimulations to electrode 1 (E1) are then delivered, followed by a 30-second rest, then 12 trains of stimulations to electrode 2 (E2). Dopamine [10 μM] is perfused in the chamber for 60 seconds paired with either E1 or E2 stimulation. The brain is then rested for 15 minutes before being tested by 3 trains of E2 stimulation followed by 3 trains of E1 stimulation (post-induction). ***G,*** Representative trace of a prep in which dopamine perfusion is paired with E1 stimulations. ***H,*** Mean relative change of the calcium responses in the α’3 compartment. Mean ±SEM. Unpaired t-test, two tailed; n = 8: ***t_(14)_ = 4.505, p = 0.0005.

In pre-induction and post-induction AL stimulations, 5 trains of 20 pulses at 100Hz were applied. Pulse width is 1 millisecond and inter-pulse interval is 9 milliseconds. Inter-train interval is 15 seconds. Stimulation strength is 100 (low input stimulation) or 200 (high input stimulation) μAmps. During induction, 12 AL stimulation trains were applied with 5 seconds inter-train interval. Regarding the AL electrode size, we noticed a relationship between the diameter of the AL electrode tip and the input current such that applying a 100 μAmps input stimulation via a large electrode tip diameter has a similar effect on baseline KCs calcium responses and on the resultant plasticity as applying a 200 μAmps via a small electrode tip diameter. For example, with a 100 μAmps input stimulation, PDP can be formed in the MB α3 compartment if the diameter of the AL electrode is significantly enlarged. Therefore, to minimize variability within the same dataset, all electrodes used in any experiment in this study were made at the same time using a p-97 micropipette puller (Sutter Instruments) before data collection. To minimize variability across the different experiments, we kept the AL electrode size approximately one fourth of the AL size.

AFV stimulation was similar to AL stimulation but the diameter of the AFV stimulation electrode was large enough to suck in the free end of the AFV. In GRAB_DA2m_ experiments, AFV stimulation strength was adjusted to be above the threshold of KC GRAB_DA2m_ responses; it varied between 500 to 1000 μAmps. We noted that AFV stimulation under these conditions did not produce a GRAB_DA2m_ response in horizontal lobes, suggesting that while PPL1 neurons were stimulated, PAM neurons were not. Whether this reflects differences in circuitry or differences in relative excitability (note we are using HL3) is unknown. In AL pairing experiments, we noted in tests of the AFV electrode that there were MB GCaMP responses in some animals, but not all (2/8 had no response). In no case, however, did AFV stimulation alone cause PDP suggesting that AFV-stimulated MB calcium increases are not able to act as a CS. Plasticity has been observed to be dependent on calcium entry pathway previously in mammalian brain (Deisseroth et al., 1998).

Images were captured using an ORCA Flash4.0 V3 sCMOS camera at 200 frames per second (except in the MBON imaging experiment using *MB027B split-GAL4>20x-UAS-GCaMP6f*, frame rate was reduced to 2 frames per second) with a 40x water immersion lens on an Olympus upright microscope BX50W1. Images were collected as 512x512 resolution and a binning factor of 2x2. Imaging was done using the HCImage Live software. Excitation of the used florescent sensors was done using the CoolLED pE-4000 LED source. For GCaMP and GRAB_DA2m_, the 470 nm LED channel was used with an excitation filter Chroma 450/50 and emission filter FF01-525/45-25. jRCaMP excitation was done using the 550 nm LED channel and the excitation filter FF01-530/43-25 and an emission filter FF01-607/36-25. Calcium traces at every frame were calculated as ΔF/F_0_ = (F-F_0_)/F_0_ where F is the florescence value at a given frame and F_0_ is the florescence value at baseline. Peak response to a stimulation was calculated by subtracting the average ΔF/F_0_ during the last second before a stimulation from the peak ΔF/F_0_ during stimulation. The responses to AL stimulations before and 15 minutes after induction were averaged to calculate the pre and post responses, respectively. PDP values or ΔF/F relative change was calculated as (post-pre)/pre.

The isolated pulse stimulator model 2100 and the perfusion system ValveLink 8.2 were triggered using a custom program written and controlled by the pClamp 11 software and the Digidata1550A digitizer.

### Behavioral experiments

Aversive learning experiments were performed in an environmental room in red light at 25°C with 65% humidity. Flies (mixed males and females) were between 4-14 days old. Flies were given at least 10 minutes acclimation period in room before training or testing. Data for each experiment was pooled from at least 3 independent experimental days. The learning assays were performed as described by the Quinn lab (Tully and Quinn, 1985). The US was provided as 12 1-second 90- or 60-volts shocks during the 1-min CS-US pairing. 10% 4-methylcyclohexanol (MCH) and 3-octanol OCT were used as the CS odors. Flies were then given a 2-minute rest. Testing involved 2 minutes of simultaneous exposure to CS odors, after which flies choosing either odor were counted. A performance index (PI) was calculated for each trial as (number of flies choosing the conditioned odor) - (number of flies choosing the not-conditioned odor) / (total number of flies). This PI was averaged between reciprocal trials where one of the odors was conditioned in one trial and the other odor was conditioned in the other to calculate the Learning Index (LI). To confirm the *CaMKII^UDel^*flies sensitivity to electric shock, 2-minute preference tests were performed during which flies chose between a stimulus vial (24 spaced 1-s 90 V shocks) and a neutral vial.

### Sleep Assay

Sleep deprivation was done in 25°C incubators on a 12 h light, 12 h dark cycles. Mated female flies were individually loaded into glass sleep tubes containing a food mixture of 5% sucrose and 2% agar. Drosophila Activity monitors (DAM) system (TriKinetics, Waltham) was used to measure sleep. Sleep was defined as inactivity bouts of 5 or more minutes (Hendricks et al., 2000; Shaw et al., 2000). Flies were sleep deprived by turning on the shaker between ZT12-ZT24. DAM data was analyzed using a custom MATLAB program (Donelson et al., 2012).

### Immunostaining

Adult fly brains were dissected in cold Schneider’s Insect Medium (Sigma, S0146), and then fixed in 4% PFA solution for 30 mins at room temperature. Fixed brains were washed 3x30 mins in 0.5% Triton-PBS (PBST) solution, blocked in 10% normal goat serum solution for 1 hour, and incubated in mouse anti-CaMKII antibody (1:10,000, Cosmo) for 3 days. CaMKII antibody solutions were removed, and samples washed in PBST solution for 3x30 mins. Samples were then incubated in Alexa Fluor 633 anti-mouse antibody (Invitrogen) overnight, then washed 3x30 mins in PBST solution and mounted in the Vectashield mounting medium. Images were taken under a 20x objective lens with the same settings using a Leica SP5 confocal microscope. The images were analyzed by ImageJ software. For the intensity of MB regions, the middle slices were selected as the representative pictures, and mean intensity of all MB lobes were quantified.

### Immunoblotting

100 5-day old *Canton-S* wildtype (WT) or *CaMKII^UDel^* flies (mixed males and females) were frozen on dry ice and vortexed to remove heads. Fly heads were separated from the fly bodies using a sieve. The heads were then homogenized in loading buffer (4X Bolt LDS, Invitrogen, Novex with 5% β-mercaptoethanol added) and heated for 10 min. Proteins were separated by SDS-PAGE (Bolt, Bis-Tris Protein Gels, Invitrogen) and transferred to a nitrocellulose membrane (GE Healthcare). Membrane was blocked (Blocking Buffer for Fluorescent Western Blotting, Rockland Immunochemicals) and then incubated with Anti-dCaMKII Clone 18 (1:1000, CosmoBio) and anti-actin mAb C4 (1:1000, Millipore). The secondary antibody used was, DyLight 680 mouse. Membrane was imaged using ChemiDoc system from Bio-Rad. Intensity of bands was calculated using Adj. Volume (Int) within the ImageLab 6.0 software. Intensity of the CaMKII band was normalized to that of actin in the same lane.

### Experimental Design and Statistical Analysis

Female flies were used in all imaging experiments due to their larger size. Both males and females aged 4 to 8 days were used in other experiments unless noted otherwise. All statistical analyses were performed in Prism 9 software. All tests were two tailed and confidence levels were set at α = 0.05. Normality of statistical data was determined via the Shapiro-Wilk test (α=0.05). Parametric tests were used for all experiments except the immunoblotting experiment where non-parametric tests were used. The statistical tests, p-values, sample sizes and other statistical information for each experiment are listed in figure legends. Post-hoc analysis information for Figure 2G and Figure 5F are listed in Table 1 and Table 2.

**Table 1.** Tukey’s post-hoc results for Figure 2G. Table shows statistical data for each comparison in Figure 2G in □’3 and □’3 compartments. N1, sample size of first group; N2, sample size of second group; q, value of studentized range distribution; DF, degrees of freedom; p, p-value.

**Table 2.** Šídák post-hoc results for Figure 5F. Table shows statistical data for each comparison in Figure 5F. N1, sample size of first group; N2, sample size of second group; t, value of the t-distribution; DF, degrees of freedom; p, p-value.

## Results

### Artificial aversive training induces a potentiation PDP in KCs

Previous studies had demonstrated that enhanced calcium responses similar to those reported in the MB *in vivo* after aversive training can be achieved in dissected brains (Wang et al., 2008; Tomchik and Davis, 2009; Ueno et al., 2013). To investigate whether this plasticity recapitulates the formation of aversive memory, and to optimize our protocol, we dissected the central nervous system (brain with attached ventral nerve cord) of 4-8 days old mated female flies expressing the Ca^2+^ indicator GCaMP6f in the MBs using the KCs driver (*VT030559*-*GAL4*). Dissected brains were pinned in an imaging bath chamber and the cervical connective towards the base of the ventral nerve cord was cut to allow electrical stimulation of the ascending fibers of the ventral nerve cord (AFV) connecting the ventral nerve cord to the brain (Figure 1A). We used glass suction electrodes to stimulate the antennal lobe projection neurons and the AFV to activate the odor pathway and the electric shock pathway, respectively (Figure 1B). Given that *in vivo* aversive STM formation is correlated with enhanced calcium responses in the α’3 MB compartment but not in the α3 compartment (Krashes et al., 2007; Wang et al., 2008; Cervantes-Sandoval et al., 2013; Zhang et al., 2019), we decided to focus our calcium imaging on these two regions (Figure 1C). We found that both α’3 and α3 compartments respond only to ipsilateral AL stimulation, but AFV stimulation could generate a calcium response in both compartments of both MBs. Therefore, we imaged the MB ipsilateral to the stimulated AL.

To induce the formation of an aversive STM trace, we paired 12 trains of stimulation of both the AL electrode and the AFV electrode, thus activating the CS and the US pathways at the same time (paired induction). As controls, we repeated the same induction paradigm but separated the AL stimulation from the AFV stimulation by 30 seconds (unpaired induction) or omitted the AFV stimulation (AL alone induction) or omitted the AL stimulation (AFV alone induction) or simply allowed the brain to rest for 15 mins (no induction). We compared the change in calcium response to AL stimulation before induction and 15 mins post induction (Figure 1D). We predicted that if this *ex vivo* model truly recapitulated short-term aversive training, only the paired induction should produce an enhancement of the calcium responses in the α’3 and none of the stimuli should potentiate α3 (Krashes et al., 2007; Wang et al., 2008; Davis, 2011; Cervantes-Sandoval et al., 2013). Indeed, this was the result we obtained (Figure 1E,F). Interestingly, we noticed that the repetitive unpaired activation of KCs results in suppression of their calcium responses (Figure 1F). This is reminiscent of desensitization or habituation; a non-associative plasticity/memory described as a decrement in neural responses to uninteresting, frequently-encountered stimuli whose predictive value is negligible (Wilson and Linster, 2008; Wilson, 2009). These non-associative processes have been documented in insects (Cho et al., 2004; Das et al., 2011; Semelidou et al., 2018), rodents (Wilson, 1998; Best and Wilson, 2004), and humans (Ferdenzi et al., 2014; Pellegrino et al., 2017).

### Dopamine replaces the artificial US stimulus but does not replace the CS

Multiple behavioral and *in vivo* imaging studies have shown that dopaminergic neurons encode the US valence information in the MB, with PPL1s providing aversive reinforcement and PAMs providing appetitive reinforcement (Schwaerzel et al., 2003; Riemensperger et al., 2005; Schroll et al., 2006; Kim et al., 2007; Claridge-Chang et al., 2009; Mao and Davis, 2009; Aso et al., 2010; Aso et al., 2012; Pech et al., 2013; Aso et al., 2014b; Yamagata et al., 2016; Handler et al., 2019). These studies suggest that the US alone is sufficient to evoke dopamine release. However, mechanistic studies employing the LTE paradigm reported no dopamine release after US pathway stimulation alone. Strong dopamine release was only seen after coincident activation of both the CS and the US pathways, and it was concluded that dopamine release is downstream of the CS+US coincidence and does not encode the primary US information (Ueno et al., 2017; Ueno et al., 2020). To directly address this discrepancy, we expressed a G-protein-coupled receptor-activation-based dopamine sensor (*GRAB_DA_*) (Sun et al., 2020) in the KCs. We then dissected the fly’s central nervous system and activated the US pathway by stimulating the AFV. We also expressed the calcium indicator jRCaMP1a in the KCs to use as a reference for the strength of activation (Figure 2A,B). We observed a robust dopamine release in the α’3 and a much weaker release onto the α3 compartment in response to the same AFV stimulation used in our induction experiments (Figure 2C,D). This result shows that dopamine release occurs in response to the US stimulus alone and does not require CS+US coincidence.

We next asked if dopamine perfusion can replace electrical stimulation of the AFV. We paired electrical stimulation of the AL with perfusion of different concentrations (1, 5 and 10 μM) dopamine and analyzed the KC response to AL stimulation before and after the pairing (Figure 2E,F). Similar to the plasticity seen after AL+AFV induction, we found that only the paired AL+ DA (5 or 10μM) induction induced an enhancement PDP in the α’3; pairing with 1 μM DA was not sufficient to induce PDP (Figure 2G; detailed statistical data for the post-hoc analysis are listed in Table 1). Also, in agreement with our observation that dopamine release in α3 was very weak compared to the α’3 compartment upon AFV stimulation, our protocol did not induce *ex vivo* PDP in the α3 compartment (Figure 2G). We also found that PDP (induced by AL + 10 μM DA) lasts for at least one hour after induction (Figure 2H).

Although these results supported a PDP model with direct dopamine modulation of KCs, we were concerned with the lack of synaptic specificity in the AL+DA induction, as dopamine was perfused onto the whole preparation. To address this concern, we repeated the experiment but replaced dopamine perfusion with activation of the dopaminergic neurons in the PPL1 cluster, which carry the negative valence US information to the MB (Schroll et al., 2006; Claridge-Chang et al., 2009; Mao and Davis, 2009; Aso et al., 2010; Aso et al., 2012; Burke et al., 2012; Aso and Rubin, 2016). We expressed the ATP-gated channel *P2X_2_* (Yao et al., 2012) in the PPL1 dopaminergic cluster using the *TH-LexA* driver. We paired AL stimulation with application of 2.5 mM ATP to activate PPL1s (Figure 2I,J). Again, we observed an enhancement PDP in the α’3 but not in the α3 compartment (Figure 2K).

Taken together, our findings agree with previous behavioral studies and *in vivo* imaging studies and show that dopamine is released in response to the US alone and carries US valence information to the MB to allow associative learning. Potentiation PDP induced in our *ex vivo* paradigm also localizes to α’ cells only, congruent with the memory traces reported *in vivo* with aversive short-term memory.

### PDP *ex vivo* plasticity is input specific

Associative memory is CS-specific. A fly that is aversively trained against an odor shows aversive behavior to that odor and closely similar odors only, but does not generalize the aversion to all odors (Barth et al., 2014). Olfactory information is sparsely transmitted to KC dendrites, with the identity of the odor being determined by population coding (Marin et al., 2002; Perez-Orive et al., 2002; Wong et al., 2002; Murthy et al., 2008; Turner et al., 2008; Honegger et al., 2011). To mimic the application of 2 different odors to the fly brain, we placed two suction glass electrodes onto the same antennal lobe and spaced them as far as possible from each other to maximize the probability that each electrode stimulates a distinct subset of projection neurons. Theoretically, a certain degree of overlap between the two subsets could be allowed as long as the two activated populations are sufficiently distinct. To test if the two electrodes activated distinct populations of projection neurons, we expressed the calcium indicator (GCaMP6f) in the projection neurons and divided the antennal lobe into multiple regions of interest asking if the pattern of responses were equivalent for the two electrodes (Figure 3A,B). We found that spacing resulted in activation of distinct populations of projection neurons, allowing us to provide two hypothetical odors to the MB (Figure 3C,D).

We then asked whether the observed *ex vivo* PDP is specific to the olfactory input activated during induction. To challenge the olfactory-input specificity of our preparation even more, we used the less specific induction method, AL+DA, in which dopamine is impartially perfused to the whole brain to eliminate any specificity on the US pathway side (Figure 3E). We recorded KC calcium responses to both electrode stimulations before induction (pre). Then, we delivered a train of stimulations through electrode #1, followed by a train of stimulations through electrode #2. Dopamine perfusion started 5 seconds before the stimulation train for only one of the two electrodes (paired) and stopped at the last pulse of the train, while normal HL3 saline was perfused during the activation of the other electrode (unpaired). We then allowed the brain to rest for 15 mins and tested the KC response to electrode #2 first then to electrode #1 to eliminate any bias due to the order of stimulation (Figure 3F). An increased calcium response was observed in α’3 after activation of the paired subset compared to the unpaired one (Figure 3G,H). This result demonstrates that even with the broader bath-dopamine induction paradigm, the plasticity achieved in our paradigm is specific to the CS input activated during *ex vivo* training.

### Artificial aversive training induces a suppression PDP in α’3 MBONs

While the α’ branches of the KCs are potentiated in response to an aversively trained odor in intact animals, responses in their postsynaptic MBONs are suppressed (Sejourne et al., 2011; Owald et al., 2015; Owald and Waddell, 2015; Zhang et al., 2019). To eliminate the possibility that our paradigm potentiates memory-relevant KCs via a memory-irrelevant epiphenomenon, we needed to demonstrate that our *ex vivo* training could produce a suppression PDP in MBONs. Therefore, we repeated the same AL+DA induction described previously (Figure 2F), but this time we expressed *20xGCaMP6f* with the *MB027b split-GAL4* driver to examine the responses in the α’3 MBONs (MBON-α′3ap and MBON-α′3m) (Tanaka et al., 2008; Sejourne et al., 2011; Aso et al., 2014a) (Figure 4A). We found that the α’3 MBON response to AL stimulation is suppressed after our artificial learning paradigm (Figure 4B,C), agreeing with *in vivo* results. These findings show that both the enhancement and suppression PDPs achieved in our paradigm are learning-specific and are encoded as enhancement in the presynaptic KCs and a suppression in the postsynaptic MBONs.

**Figure 4.**
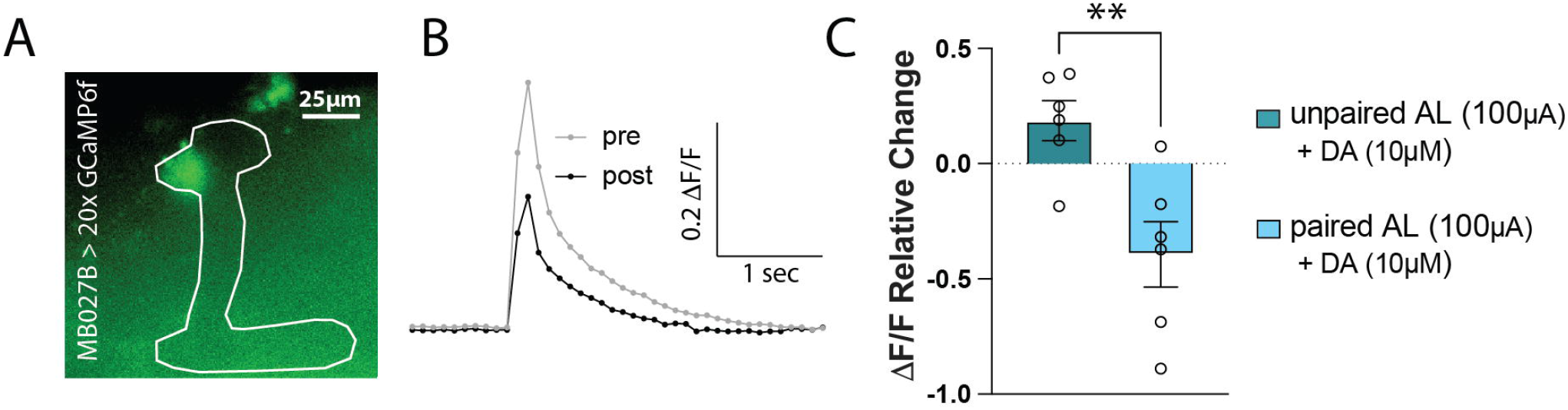
α’3 MBONs dendrites show a suppression PDP. ***A,*** Representative image of the GCaMP6f signal expressed in the α’3 MBONs using the *MB027B* splitGAL4 line. The MB is outlined in white, showing the localization of the analyzed dendritic signal in the α’3 compartment. Scale bar is 25 μm. ***B,*** Representative trace of the α’3 MBONs responses pre and post the AL+DA induction as described in Figure 2F. ***C,*** Mean relative change of the calcium responses in the MBONs’ dendrites in the α’3 compartment. Mean ±SEM. Unpaired t-test, two tailed; n = 6: **t_(10)_ = 3.495, p = 0.0058.

### The 3’UTR mRNA of *CaMKII* is important for *ex vivo* PDP

We next turned our attention to exploring the utility of this paradigm for understanding memory formation, choosing problems at several different levels of analysis: molecular components of the memory machinery, organizational principles of the circuitry and the role of brain state in gating plasticity. At the molecular level, previous studies have demonstrated that CaMKII is important for synaptic plasticity and memory formation in many species (Kelleher et al., 2004; Giese and Mizuno, 2013) including *Drosophila melanogaster* (Griffith et al., 1993; Koh et al., 1999; Ashraf et al., 2006; Malik et al., 2013; Mitchell et al., 2021). Recently, we found that the long 3’UTR region of *CaMKII* mRNA is responsible for the activity-dependent synthesis of CaMKII in presynaptic terminals at the larval neuromuscular junction (Kuklin et al., 2017) and for the basal accumulation of axonal CaMKII protein in MB (Chen et al., 2022). We hypothesized that the loss of the 3’UTR would impair associative plasticity by decreasing synaptic CaMKII protein levels and disrupting the signaling machinery that is triggered by the CS+US coincidence. We tested this idea and used our *ex vivo* PDP preparation to gain insight into whether this effect is upstream or downstream of the CS+US coincidence.

First, we asked if loss of the 3’UTR of *CaMKII* mRNA affects CaMKII levels. We used animals in which the *CaMKII* gene was engineered using CRISPR/Cas9 to lack the 3’UTR (*CaMKII^Udel^*, Chen et al., 2022) (Figure 5A). Immunostaining to quantify CaMKII protein levels in the MB neuropil in both wildtype and *CaMKII^UDel^*showed that CaMKII levels are specifically decreased in synaptic regions (Figure 5B). We also used western blotting to quantify total CaMKII levels and found a substantial decrease, normalized to actin, in *CaMKII^UDel^* flies compared to wildtype (Figure 5C). This is in agreement with a previous study which found that mice expressing a mutant form of CaMKII lacking the 3’UTR show decreased levels of CaMKII in the dendritic, but not the somatic, region of hippocampal neurons (Miller et al., 2002).

**Figure 5.**
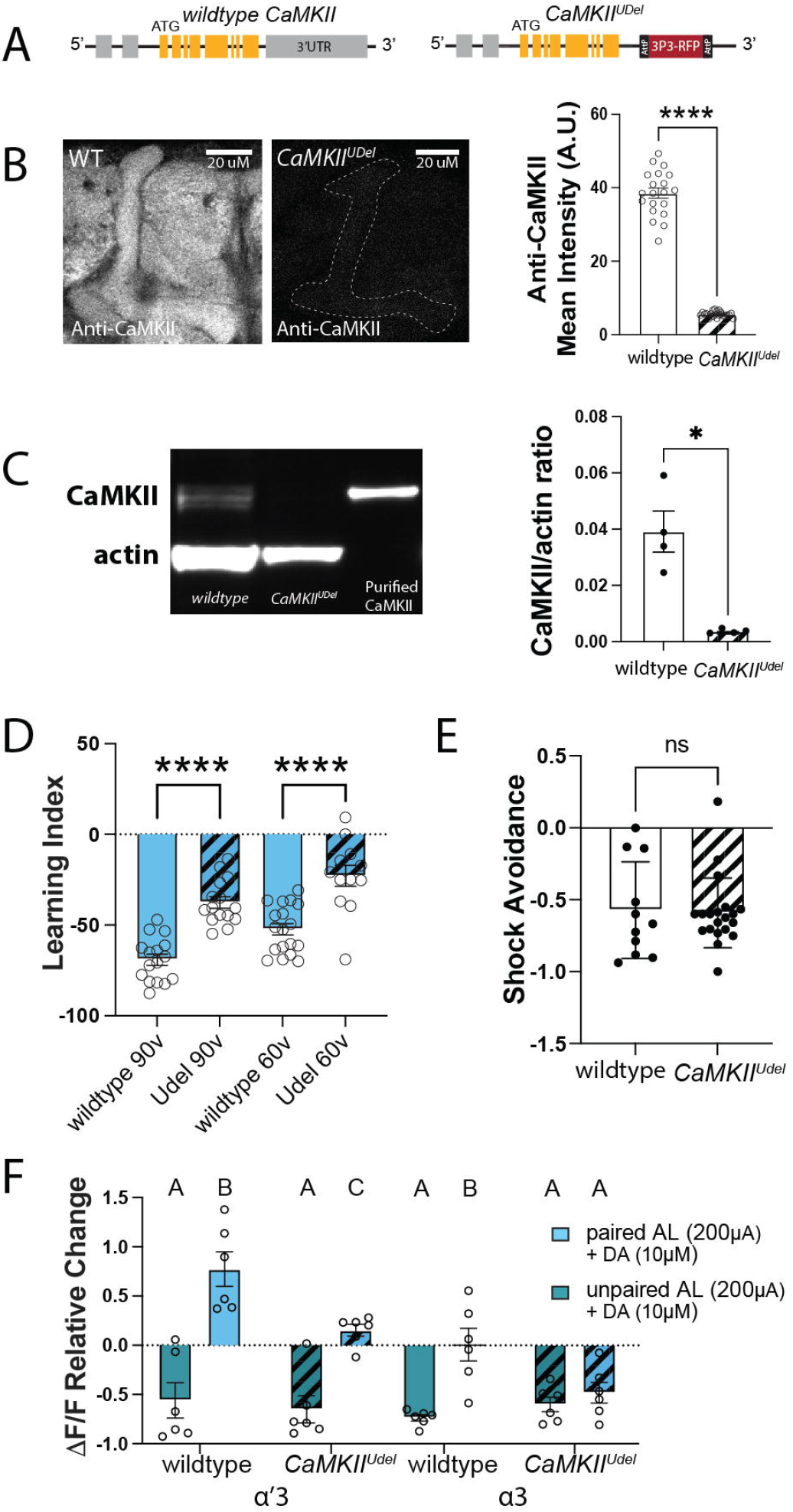
The 3’UTR region of *CaMKII* is important for PDP formation. ***A,*** Schematic of wildtype (left) and *CaMKII^Udel^* an allele in which the 3’UTR has been replaced with an RFP marker using CRISPR/Cas9 (right). ***B,*** (Left) Representative immunostaining images showing CaMKII protein levels in the MB in wildtype *CaMKII* flies and in *CaMKII^Udel^* flies. Dotted line shows position of the MB. Scale bar is 20 μm. (Right) quantification of CaMKII levels, mean±SEM. Unpaired t-test, two tailed; n = 20 wildtype flies and 22 *CaMKII^Udel^*flies: ****t_(40)_ = 24.81, p < 0.0001. ***C,*** (Left) Western blot showing CaMKII and actin levels in wildtype or *CaMKII^Udel^*adult female brains and in purified CaMKII samples. (Right) Quantification of CaMKII immunoreactivity mean ±SEM. Mann-Whitney test; n = 4 in each condition; sum of ranks in wildtype group is 26; sum of ranks in *CaMKII^Udel^* group is 10: *p=0.0286.***D,*** Learning index of wildtype and *CaMKII^Udel^*flies after training with 90 volts or 60 volts. Mean ±SEM. Two-way ANOVA: punishment voltage effects F_(1, 57)_ = 18, p<0.0001; 3’UTR effects F_(1,57)_= 67.57, p<0.0001. Šídák post-hoc tests: when punishment is 60 volts, ****t_(57)_ = 5.516, p<0.0001; when punishment is 90 volts, ****t_(57)_ = 6.122, p<0.0001. ***E,*** Wildtype and *CaMKII^Udel^* flies can avoid electric shock. Mean ±SEM. Unpaired t-test; data collected across 11 trials of the wildtype group and 20 trials of the *CaMKII^Udel^*group; number of flies per group ranged between 21 to 47 with an average of 29 flies per group: ^ns^p=0.8518. ***F,*** Mean relative change of the calcium responses in the α’3 and the α3 compartments after unpaired AL+DA induction (green) and paired AL+DA induction (blue) in wildtype flies or *CaMKII^Udel^*flies. Mean ±SEM. Three-way ANOVA; n=6 in each condition: lobe effects F_(1,40)_ = 17.52, p= 0.0002; induction effects F_(1,40_) = 68.43, p<0.0001; 3’UTR effects F_(1,40)_ = 8.625, p=0.0055; 3’UTR x lobe interaction effects F_(1,40)_ = 0.9631, p=0.3323; 3’UTR x induction interaction effects F_(1,40)_ = 10.16, p = 0.0028; lobe x induction effects F_(1,40)_ = 12.25, p = 0.0012; 3’UTR x lobe x induction interaction effects F_(1,40)_ = 0.07388, p =0.7872. Šídák post-hoc results are listed in Table 2. Stripe bars indicate the *CaMKII^Udel^* genotype.

We then tested the impact of the 3’UTR deletion on aversive STM. We found that *CaMKII^UDel^* flies showed a significant impairment in immediate memory performance compared to wildtype flies (Figure 5D). This impairment was not due to sensorimotor dysfunctionalities as both wildtype and *CaMKII^UDel^*flies respond to electric shock (Figure 5E). We then tested the effect of the *CaMKII* mRNA 3’UTR deletion on potentiation PDP in KCs. We used the previously described AL+DA induction protocol and examined KC responses to AL stimulation both before and 15 mins after either paired or unpaired induction. In the α’3 compartment, although both wildtype and *CaMKII^UDel^* genotypes showed statistically significant PDP relative to the unpaired induction, the PDP in the wildtype flies was more than 5 times stronger than in the *CaMKII^UDel^*flies (Figure 5F; detailed statistical data for the post-hoc analysis are listed Table 2). We also noticed that the change in signal in the α3 compartment in wildtype flies after paired induction was statistically different from the unpaired induction. However, this does not translate into formation of PDP in that compartment as this change was not different from zero (one-sample t-test, p-value 0.947). This is likely due to a technical difference as the AL input stimulation in this experiment was stronger (200 μAmps vs. 100 μAmp in previous experiments) which led to a stronger suppression after the unpaired AL stimulation in the unpaired induction (see also below). These results show that the 3’UTR mRNA of *CaMKII* is important for memory and its loss impairs learning-induced plasticity. It also indicates that the effect on the learning circuit is downstream of the CS+US coincidence and does not block the CS+US coincidence detection machinery itself.

### Stronger inputs to the MB can recruit the α3 compartment into the short-term memory circuit

At the circuit level, it has been hypothesized that memory in flies, like humans (McClelland et al., 1995; Dudai, 2012), undergoes systems consolidation: initial potentiation of the α’β’ lobes with a time-dependent transfer of potentiation to αβ. The MB α’3 and α3 compartments are adjacent to each other. Both respond to odor (Turner et al., 2008) and AL stimulation (Figure 6B), and both use the same coincidence detector, *rutabaga* (Livingstone et al., 1984; Levin et al., 1992; Mao et al., 2004; Gervasi et al., 2010). This raises the question of how the two compartments are able to play different roles in the memory circuit and why only the α’β’ cells show immediate PDP both *ex vivo* and *in vivo*. To determine if this might be due to differences in the intrinsic properties of the two classes of cells, we examined the basal responses of both compartments to a ramp of AL stimulations without any pairings (Figure 6A). We found that α’3 axons are recruited first, with very weak AL stimulation, while α3 only starts responding at higher stimulation strengths (Figure 6B). There was also a distinct difference in the maximum response levels: α’3 compartment responses plateau at a much lower level than α3 (Figure 6B), likely explained by the fact that the α3 compartment receives more axons than α’3 (Li et al., 2020). Previous studies reported that odors activate a small percentage of KCs (∼5-12%), and that the odor responses in α’β’ cells are stronger than those of αβ cells (Turner et al., 2008; Inada et al., 2017). This suggests that actual odor encoding is similar to the weak AL stimulation used in our experiments (100 μAmp), recruiting only α’3 but insufficient to recruit α3. We hypothesized that an increase in the AL stimulation intensity during *ex vivo* induction might be sufficient to recruit α3 and allow PDP in that compartment as well. We repeated both AL+AFV and AL+DA inductions, but this time with a 200 μAmp AL stimulation intensity. Indeed, under these conditions we found that PDP occurred in both the α’3 and α3 compartments (Figure 6C,D).

**Figure 6.**
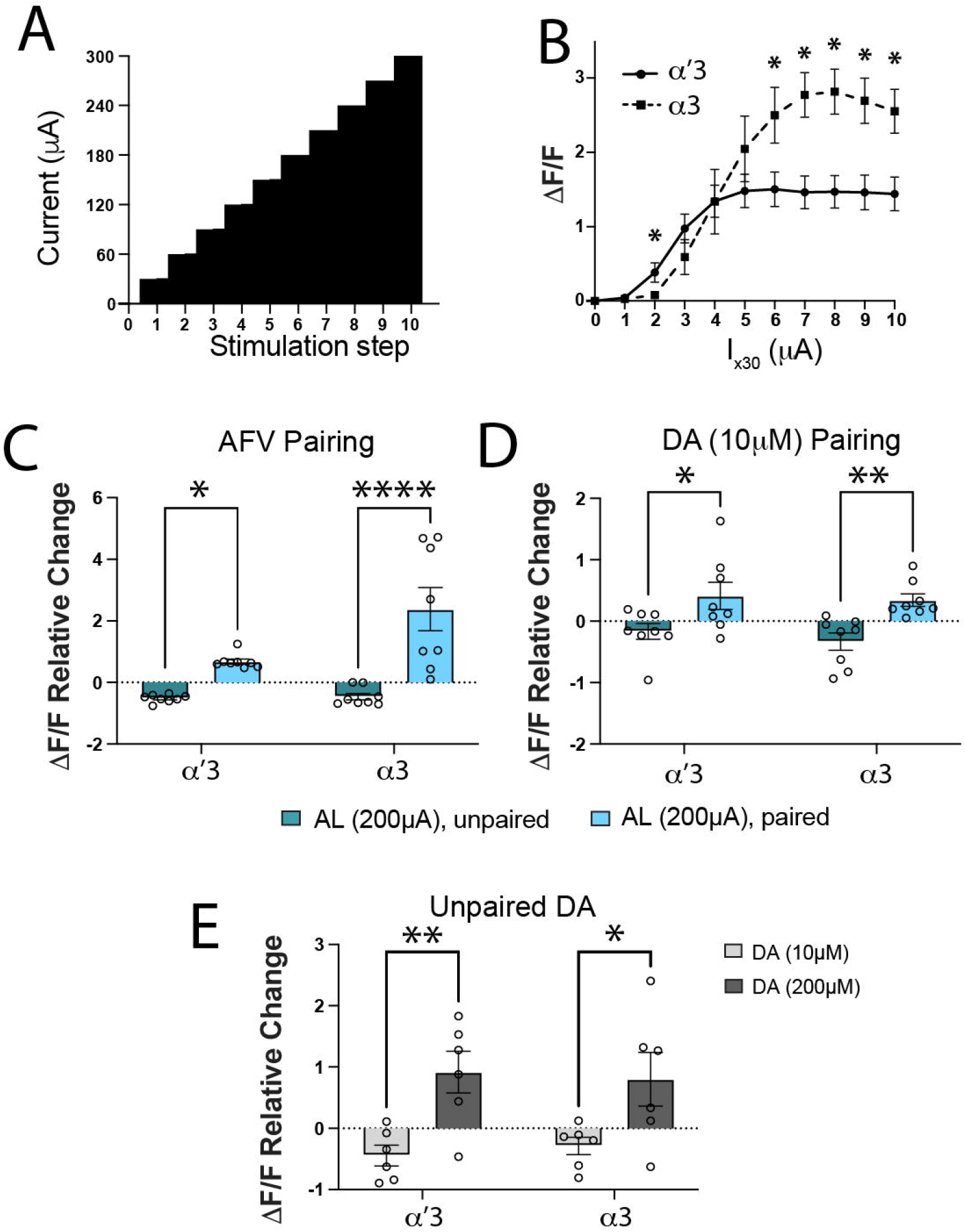
Increasing CS or US pathway activation recruits the α3 compartment to the PDP circuit. ***A,*** Design of the AL ramp. AL input stimulation is increased by 30 μAmps every 30 seconds. ***B,*** Unpaired calcium responses to AL stimulations in the α’3 and α3 compartments showing mean±SEM. Unpaired t-test; n=8 *p<0.05: t_(14)_ = {1.873, p=0.082; 2.304, p=0.0371; 1.254, p=0.23; 0.00567, p=0.99555; 1.143, p=0.2722; 2.267, p=0.0397; 3.517, p=0.00342; 3.6, p=0.002898; 3.218, p=0.006193; 2.979, p=0.009959} for comparisons between α’3 and α3 responses to {30; 60; 90; 120; 150; 180; 210; 240; 270; 300} μAmps AL stimulation, respectively. ***C and D,*** Mean relative change of the calcium responses in the α’3 and the α3 compartments after the unpaired (green) or paired (blue) AL+AFV induction in ***C*** and AL+DA induction in ***D*** when the AL input stimulation is increased to 200 μAmps. Mean ±SEM. In ***C***, Two-way ANOVA (α = 0.05; n = 8 in each condition): lobe effects F_(1, 28)_ = 6.078, p= 0.0201; induction effects F_(1, 28)_ = 32.08, p<0.0001; lobe x induction interaction effects F_(1,28)_ = 5.338, p=0.0285. Šídák post-hoc tests: In the α’3 compartment, *t_(28)_ = 2.372, p=0.049; In the α3 compartment, ****t_(28)_ = 5.639, p<0.0001. In ***D***, Two-way ANOVA (α = 0.05; n = 8 in each condition): lobe effects F_(1, 28)_ = 0.5793, p= 0.4530; induction effects F_(1, 28)_ = 16.36, p=0.003; lobe x induction interaction effects F_(1,28)_ = 0.1038, p=0.74985. Šídák post-hoc tests: In the α’3 compartment, *t_(28)_ = 2.656, p=0.0256; In the α3 compartment, **t_(28)_ = 3.112, p=0.0085. ***E,*** Mean relative change of the calcium responses in the α’3 and the α3 compartments after unpaired 10μM or 200μM DA perfusion alone, stimulation at 100 μAmps. Two-way ANOVA (α = 0.05; n = 6 in each condition): lobe effects F_(1, 20)_ = 0.005471, p= 0.9418; concentration effects F_(1, 20)_ = 16.88, p=0.0005; lobe x concentration interaction effects F_(1,20)_ = 0.2026, p=0.6574. Šídák post-hoc tests: In the α’3 compartment, **t_(20)_ = 3.223, p=0.0085; In the α3 compartment, *t_(20)_ = 2.586, p=0.035.

This result agrees with predictions from our previously published theoretical model of the associative learning circuit in which the CS+US coincidence triggers a recurrent loop between KCs and dopaminergic neurons which increases dopamine release only onto cells receiving both inputs together (Adel and Griffith, 2021). The weak response of the α3 compartment to weak AL stimulation is subthreshold for the MB coincidence detector, and insufficient for forming PDP in α3. Stronger AL stimulation puts α3 over threshold and triggers the coincidence detector in α axons, thus increasing dopamine gain and forming PDP/memory.

Our model further predicts that there should be a certain dopamine concentration above which dopamine alone is sufficient to potentiate both α’3 and α3 compartments without any need for pairing, as this hypothetical high dopamine concentration bypasses the need for CS+US coincidence detector activation of the gain control machinery that increases local dopamine release. To test this prediction, we applied either 10 μM dopamine (used in our previous experiments) or a high dopamine concentration (200 μM) to the dissected brains without pairing with AL stimulation. Indeed, 200 μM dopamine alone was sufficient to potentiate both α’3 and α3 responses to AL stimulation (100 μAmps) 15 minutes post dopamine application (Figure 6E). A similar effect of high dopamine was reported by Ueno et al., 2013, but interpreted differently. We show that pairing-independent plasticity resulting from very high dopamine, while possibly implemented by the same machinery responsible for associative learning, does not truly represent a memory trace as it lacks specificity to CS-activated synapses, likely representing a generalized potentiated state of the MB. In contrast, the PDP seen with lower dopamine concentrations relies on an interplay between the CS and US signals to gate dopamine gain only at the synapses activated by the CS during training.

### PDP is blocked by sleep deprivation and rescued by rebound sleep

Many of the most interesting questions about memory formation are ones that involve the interaction of plasticity with the animal’s internal state. To test the ability of this preparation to retain traces of previous experience and allow interrogation of memory mechanisms at this level, we looked at the effect of sleep on subsequent *ex vivo* plasticity. Sleep is linked to memory across phyla. In *Drosophila*, perturbations of sleep have been shown to impair both memory formation (Bushey et al., 2007; Seugnet et al., 2008; Li et al., 2009; Seugnet et al., 2009; Donlea et al., 2014; Seidner et al., 2015) and consolidation (Ganguly-Fitzgerald et al., 2006; Donlea et al., 2009; Bushey et al., 2011; Dag et al., 2019); reviewed in (Goel et al., 2009; Diekelmann and Born, 2010; Dissel et al., 2015; Donlea, 2019). To determine whether the plasticity we observe in the MB is sleep-dependent, we mechanically sleep deprived entrained (12:12 light:dark) flies for 12 hours during the ZT12-24 night period before using the AL+DA paradigm to induce the artificial memory at ZT0. Control flies were housed on a different shaker in the same incubator, but the shaker was turned off. A third group of flies received shaking for only the last 2 hours of the night (ZT22-24) to control for the acute physical stress that may result from the shaker. Figure 7A shows a schematic of our experimental design and Figure 7B shows the minutes of sleep for each group during the relevant time windows. We found that sleep-deprived flies did not exhibit KC enhancement PDP, while both the sleep control group and the stress control group had normal PDP (Figure 7C).

**Figure 7.**
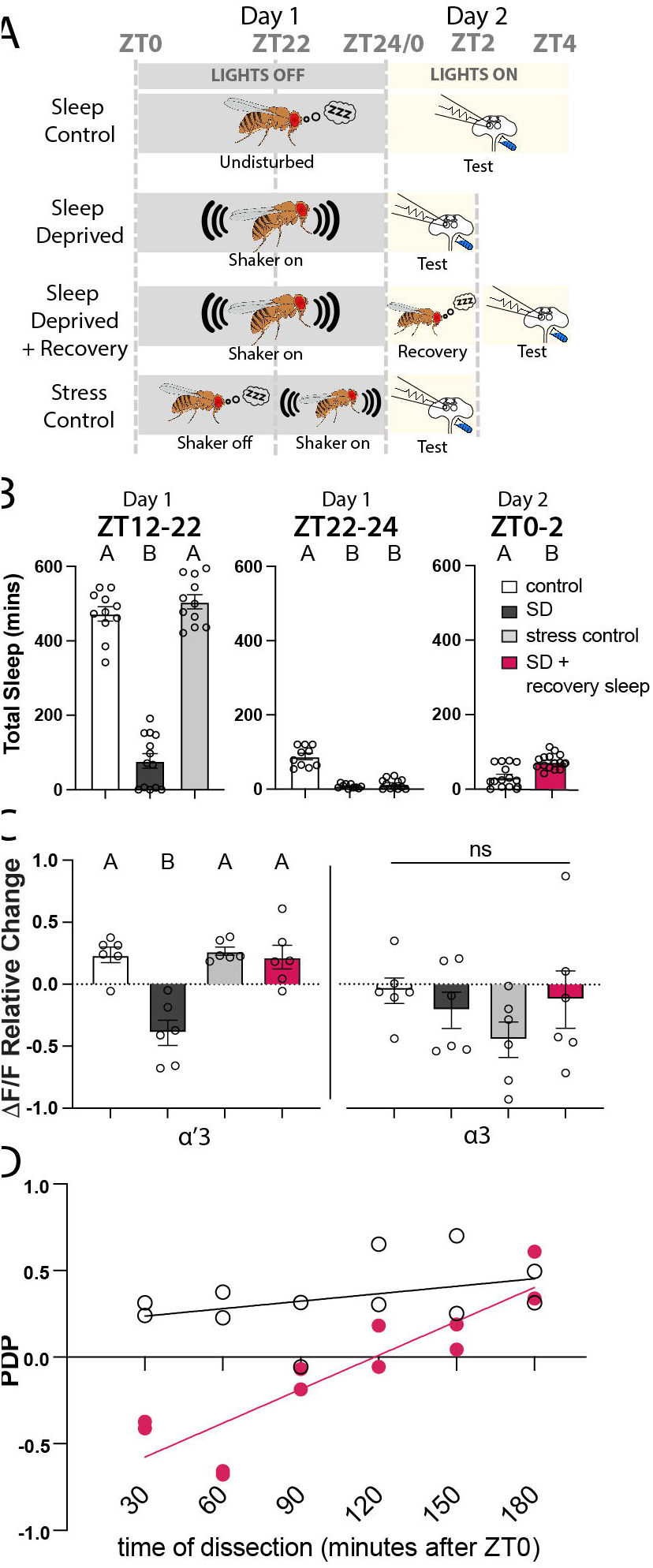
Sleep deprivation blocks PDP and rebound sleep recovers it. ***A,*** Schematic of the different sleep groups: Sleep Control, flies are undisturbed during night and tested between ZT0-ZT4 in the next day; Sleep Deprived, flies are kept on the shaker during between ZT12-ZT24 then tested between ZT0-ZT2 in the next day; Sleep Deprived + Recovery, same as Sleep Deprived but flies were allowed to recover sleep between ZT0-ZT2 then tested between ZT2-ZT4; Stress Control: flies are undisturbed during the night between ZT12-ZT22 and then the shaker was turned on between ZT22-ZT24 before testing between ZT0-ZT2 in the next day. Testing was done by the AL+DA induction described in Figure 2F. ***B,*** Total sleep of the different groups in the different ZT windows. Mean ±SEM. One-way ANOVA: in the ZT12:22 comparison, F_(2,32)_ = 159, p<0.0001; Tukey’s post-hoc {control vs. sleep deprived: q_(32)_ = 20.65, p<0.0001; control vs. stress control: q_(32)_ = 1.625, p=0.4916; sleep deprived vs. stress control: q_(32)_ = 22.34, p<0.0001). In the ZT22:24 comparison, F_(2,29)_ = 66.64, p<0.0001; Tukey’s post-hoc {control vs. sleep deprived: q_(29)_ = 14, p<0.0001; control vs. stress control: q_(29)_ = 14.47, p<0.0001; sleep deprived vs. stress control: q_(29)_ = 1.114, p=0.7136). In the ZT0:2 comparison, t_(29)_ = 4.534, p<0.0001. ***C,*** Mean relative change of the calcium responses in the α’3 (left) and the α3 (right) compartments after paired AL+DA induction. Mean ±SEM. Two-way ANOVA (α = 0.05; n = 6 in each condition): lobe effects F_(1, 40)_ = 10.10, p= 0.0029; sleep effects F_(3, 40)_ = 7.019, p=0.0007; lobe x sleep interaction effects F_(3,40)_ = 0.9308, p=0.4348. Šídák post-hoc tests: In the α’3 compartment: A vs. A, p > 0.05; B vs. A, p ≤ 0.05; t_(40)_ = {0.1592, p> 0.9999; 3.475, p= 0.0074; 0.1038, p>0.9999; 3.634, p=0.0047; 0.2631, p>0.9999; 3.371, p=0.01} for {sleep control vs. stress control; sleep control vs. sleep deprived; sleep control vs. sleep rebound; stress control vs. sleep deprived; stress control vs. sleep rebound; sleep deprived vs. sleep rebound}, respectively. No statistical significance across conditions in the α3 compartment, p > 0.05; t_(40)_ = {0.8750, p= 0.9468; 2.196, p= 0.1871; 0.4001, p= 0.9991; 1.321, p=7257; 0.475, p= 0.9977; 1.796, p=3938} for {sleep control vs. stress control; sleep control vs. sleep deprived; sleep control vs. sleep rebound; stress control vs. sleep deprived; stress control vs. sleep rebound; sleep deprived vs. sleep rebound}, respectively. ***D,*** PDP (mean relative change of dF/F) from individual animals plotted against the animal’s time of dissection. White circles indicate control animals (allowed to sleep between ZT12-ZT24) and pink circles indicate sleep deprived animals (on the shaker between ZT12-ZT24).

It is important to note that the stress control we used in this study is meant to control for acute stress, not the chronic stress that flies might experience when put on a shaker for 12 h. In many studies, the control for nighttime sleep deprivation via prolonged shaking is to sleep deprive flies during the 12 h light period. We chose not to do this for several reasons. First, it would necessitate testing the flies for PDP/memory formation at ZT12 rather than ZT0, and STM has been shown to be influenced by the clock (Lyons and Roman, 2009; Flyer-Adams et al., 2020). Second, a recent study found that sleep deprivation during the light period causes significant sleep deprivation that is discharged as rebound sleep the next day (Wiggin et al., 2020); this would invalidate using daytime shaking as a control since it produces significant sleep debt. While shaking for the last 2 hours of the night is not as stressful as 12 h of shaking, it is a better control for sleep deprivation since flies start waking up naturally in that time window, which means that there is very little sleep loss (as shown in Figure 7B).

We then asked if allowing sleep-deprived flies a period of recovery sleep could restore their ability to learn. As little as 2 hours of rebound sleep was sufficient to restore the ability to induce the same level of α’3 PDP observed in the control groups (Figure 7C). Interestingly, the 2 hours minimum of rebound sleep that we found in our experiment to be required for rescuing the *ex vivo* plasticity is similar to the period reported *in vivo* to be required for sleep-deprived flies to restore the ability to form short-term memory (Li et al., 2009). These findings demonstrate that our *ex vivo* paradigm not only forms the plasticity that is correlated with short-term memory, but also accurately recapitulates the dynamics of the learning memory circuit and its interplay with the sleep circuit. These data also suggest that the effect of sleep-deprivation on the memory circuit is downstream to sensory processing of both the CS and the US information and CS+US coincidence. This gradual recovery of the ability to form memory (Figure 7D) may support previous indications that sleep-deprivation likely impairs memory formation by downregulating dopamine receptors or other downstream molecules in the dopamine signaling pathway (Seugnet et al., 2008).

## Discussion

*Drosophila* neural circuits are traditionally studied by relating *in vivo* genetic and chemical manipulations with their consequent behavioral outcomes, from which circuit information can then be inferred (Olsen and Wilson, 2008; Simpson, 2009). More recently, the advent of *in vivo* calcium imaging allowed for tracing neural activity in actively behaving flies. Over more than a decade of such *in vivo* studies, the general circuit mechanisms of associative memory have been discovered, but there are limitations imposed by imaging the brain of an active intact fly (for review, see Adel and Griffith, 2021). These include the relatively low signal to noise ratio, the inaccessibility of multiple brain regions due to restrictions on imaging angles, the difficulty of doing acute pharmacological studies, and the possible confounds of studying the brain of a movement-restricted fly experiencing on-going stress. Taking inspiration from the way the LTP hippocampal slice model revolutionized our understanding of mammalian memory (Bliss and Lomo, 1973), we provide here an *ex vivo* model of *Drosophila* memory which can overcome these limitations and offer a powerful preparation for studying *Drosophila* memory circuits. Importantly, this model provides a framework for investigating the dynamics of neural circuits in the fly brain.

Most of the previous studies investigating the associative learning circuit *ex vivo* have focused on mapping connectivity (Cohn et al., 2015; Barnstedt et al., 2016; Felsenberg et al., 2017; Felsenberg et al., 2018; Zhao et al., 2018) or characterizing a specific biochemical pathway (Tomchik and Davis, 2009; Handler et al., 2019; Ueno et al., 2020). Only a few *ex vivo* studies (Wang et al., 2008; Ueno et al., 2013; Suzuki-Sawano et al., 2017; Ueno et al., 2017) have focused on understanding MB circuit logic. In the LTE model developed by Ueno et al (2013), pairing a stimulation of the CS and US pathways induced a potentiation of KC responses in the tips of the MB vertical lobes, but LTE did not fully recapitulate other characteristics of associative memory observed in intact flies. Here we develop a modified *ex vivo* model that resolves these discrepancies, showing that the paired activation of odor and punishment pathways induces appropriate plasticity at multiple nodes in the circuit: potentiation of KCs and suppression of MBONs. Several mechanisms for encoding those opposite forms of the plasticity have been proposed, including spike-timing-dependent-plasticity (STDP) and activation of distinct dopaminergic receptors (reviewed in Adel and Griffith, 2021). STDP mechanisms appear less likely as MBON suppression was shown to not require MBON spiking (Hige et al., 2015). Perhaps the strongest model so far comes from (Handler et al., 2019), which showed that differences in the order of KCs activation and dopaminergic input activate distinct dopaminergic receptors: DopR1or DopR2, which encode MBON suppression or potentiation, respectively. It is important to note that both (Hige et al., 2015) and (Handler et al., 2019) studied the plasticity in MB medial lobes, while our study focused on MB vertical lobes, so our paradigm may be useful in gaining greater mechanistic insight into this sign transformation in the vertical lobes.

We show that PDP is localized to the MB α’3 compartment and not in α3, in alignment with most imaging studies in intact flies. Importantly, in this *ex vivo* preparation, punishment information is relayed to the MB through dopaminergic release from the PPL1 subset. Bath application of dopamine in our preparation does not interfere with the specificity of associative learning since PDP is exclusively formed in the cells that were active during the dopamine application. These data settle several inconsistencies between previous *ex vivo* studies (Ueno et al., 2013; Ueno et al., 2017) and the majority of *in vivo* reports (Schwaerzel et al., 2003; Riemensperger et al., 2005; Schroll et al., 2006; Kim et al., 2007; Claridge-Chang et al., 2009; Mao and Davis, 2009; Aso et al., 2010; Aso et al., 2012; Pech et al., 2013; Aso et al., 2014b; Yamagata et al., 2016; Cognigni et al., 2018; Handler et al., 2019). We suggest that the genesis of the discrepancies was not due to any inherent difference between intact and *ex vivo* brains but was rather a consequence of technical considerations including stimulation strength, dopamine concentration and the sensor tools employed (see Adel and Griffith, 2021 for a more complete discussion).

An *ex vivo* preparation that recapitulates the cardinal features of the circuits underlying associative memory formation should be useful for mechanistic studies at the molecular, cellular and systems levels. We used our model to ask a new question about the innerworkings of the circuit at each of these levels. At the molecular level we demonstrated the importance of normal levels of CaMKII by manipulating the 3’UTR of *CaMKII* mRNA. Deletion of this region of the *CaMKII* gene drastically reduces the amount of CaMKII protein in synaptic regions and blunts the ability to form STM and to generate a potentiation PDP in KC axons. Our data argues that the role of this molecule is downstream of the CS+US coincidence detector, as we observe a much weaker PDP in *CaMKII^Udel^*flies. Whether the behavioral defect is due solely to the KC PDP defect is not completely clear since CaMKII likely has active roles at other circuit nodes (Mitchell et al., 2021).

At the cellular level we asked why STM and PDP form in the α’3 but not the nearby α3 compartment when both compartments respond to odors (Turner et al., 2008) and AL stimulation, and both receive dopaminergic input from the same PPL1 cluster (Masek et al., 2015). Previous work found that real odors cause activity in only 5-12% of KCs and elicit a much higher spike rate in the α’β’ KCs than in the αβ KCs. We found that low-intensity AL stimulation (100 μAmps) elicit a stronger response in the α’3 than in the α3 compartment, while high-intensity AL-stimulation (200 μAmps) causes strong responses in the α3 compartment and recruits it to the learning circuit. Coupling this with our observation of lower dopamine release in α3, suggests a model in which odor presentation during associative learning causes subthreshold responses in αβ cells such that the CS+US coincidence detector is not triggered, while the stronger responses in the α’β’ cells bypass this threshold, allowing plasticity the α’β’ cells only. This notion is in alignment with the previous finding that α’β’ cells have a lower firing threshold than αβ cells (Inada et al., 2017). Further, It is possible that long-term memory and the enhancement memory trace in the αβ KCs after repetitive space training (Yu et al., 2006) require a gradual potentiation of the αβ KCs responses with every training session such that the responses bypass the coincidence detection threshold after several training sessions. Whether repetition of AL+DA pairings recruits PDP in the α3 compartment remains unclear. It is also yet to be determined whether shortcutting the circuit and recruiting αβ cells in the first training session reduces the need for multiple spaced training sessions in long-term memory formation.

Lastly, we looked at the ability of the effects of prior experience, or brain state, on the memory circuit to be retained in the *ex vivo* preparation. Excitingly, we found that sleep deprived flies could not form PDP, but that as little as 2 hours of rest before dissection allowed the brain to recover PDP formation. The complete abolition of PDP in sleep deprived flies at first and the gradual recovery in plasticity afterwards (Figure 7D) suggests that sleep converges on the memory circuit upstream of the CS+US coincidence detector. Whether this involves regulation of dopamine receptors in the MB during sleep remains to be determined. The ability to retain in some functional way the internal state of the brain will allow this preparation to be used to understand how memory formation is altered by global system alterations.

## Acknowledgments

This work was funded by NIH R01 MH067284 to LCG. Stocks obtained from the Bloomington Drosophila Stock Center (NIH P40OD018537) were used in this study. The authors thank Dr. Kohei Ueno (Tokyo Metropolitan Institute of Medical Science) for support, encouragement, and excellent advice during the development of this project.

## References

1. Adel M, Griffith LC (2021) The Role of Dopamine in Associative Learning in Drosophila: An Updated Unified Model. Neuroscience Bulletin 37:831–852.

2. Ashraf SI, McLoon AL, Sclarsic SM, Kunes S (2006) Synaptic protein synthesis associated with memory is regulated by the RISC pathway in Drosophila. Cell 124:191–205.

3. Aso Y, Rubin GM (2016) Dopaminergic neurons write and update memories with cell-type-specific rules. Elife 5.

4. Aso Y, Siwanowicz I, Bracker L, Ito K, Kitamoto T, Tanimoto H (2010) Specific dopaminergic neurons for the formation of labile aversive memory. Curr Biol 20:1445–1451.

5. Aso Y, Herb A, Ogueta M, Siwanowicz I, Templier T, Friedrich AB, Ito K, Scholz H, Tanimoto H (2012) Three dopamine pathways induce aversive odor memories with different stability. PLoS Genet 8:e1002768.

6. Aso Y, Hattori D, Yu Y, Johnston RM, Iyer NA, Ngo TT, Dionne H, Abbott LF, Axel R, Tanimoto H, Rubin GM (2014a) The neuronal architecture of the mushroom body provides a logic for associative learning. Elife 3:e04577.

7. Aso Y et al. (2014b) Mushroom body output neurons encode valence and guide memory-based action selection in Drosophila. Elife 3:e04580.

8. Barnstedt O, Owald D, Felsenberg J, Brain R, Moszynski JP, Talbot CB, Perrat PN, Waddell S (2016) Memory-Relevant Mushroom Body Output Synapses Are Cholinergic. Neuron 89:1237–1247.

9. Barth J, Dipt S, Pech U, Hermann M, Riemensperger T, Fiala A (2014) Differential associative training enhances olfactory acuity in Drosophila melanogaster. J Neurosci 34:1819–1837.

10. Berry JA, Cervantes-Sandoval I, Chakraborty M, Davis RL (2015) Sleep Facilitates Memory by Blocking Dopamine Neuron-Mediated Forgetting. Cell 161:1656–1667.

11. Best AR, Wilson DA (2004) Coordinate synaptic mechanisms contributing to olfactory cortical adaptation. J Neurosci 24:652–660.

12. Bhandawat V, Olsen SR, Gouwens NW, Schlief ML, Wilson RI (2007) Sensory processing in the Drosophila antennal lobe increases reliability and separability of ensemble odor representations. Nat Neurosci 10:1474–1482.

13. Bliss TV, Lomo T (1973) Long-lasting potentiation of synaptic transmission in the dentate area of the anaesthetized rabbit following stimulation of the perforant path. J Physiol 232:331–356.

14. Burke CJ, Huetteroth W, Owald D, Perisse E, Krashes MJ, Das G, Gohl D, Silies M, Certel S, Waddell S (2012) Layered reward signalling through octopamine and dopamine in Drosophila. Nature 492:433–437.

15. Bushey D, Tononi G, Cirelli C (2011) Sleep and synaptic homeostasis: structural evidence in Drosophila. Science 332:1576–1581.

16. Bushey D, Huber R, Tononi G, Cirelli C (2007) Drosophila Hyperkinetic mutants have reduced sleep and impaired memory. J Neurosci 27:5384–5393.

17. Cervantes-Sandoval I, Martin-Pena A, Berry JA, Davis RL (2013) System-like consolidation of olfactory memories in Drosophila. J Neurosci 33:9846–9854.

18. Chen N, Zhang Y, Adel M, Kuklin EA, Reed ML, Mardovin JD, Bakthavachalu B, VijayRaghavan K, Ramaswami M, Griffith LC (2022) Local translation provides the asymmetric distribution of CaMKII required for associative memory formation. Curr Biol In Revision.

19. Cho W, Heberlein U, Wolf FW (2004) Habituation of an odorant-induced startle response in Drosophila. Genes Brain Behav 3:127–137.

20. Claridge-Chang A, Roorda RD, Vrontou E, Sjulson L, Li H, Hirsh J, Miesenbock G (2009) Writing memories with light-addressable reinforcement circuitry. Cell 139:405–415.

21. Cognigni P, Felsenberg J, Waddell S (2018) Do the right thing: neural network mechanisms of memory formation, expression and update in Drosophila. Curr Opin Neurobiol 49:51–58.

22. Cohn R, Morantte I, Ruta V (2015) Coordinated and Compartmentalized Neuromodulation Shapes Sensory Processing in Drosophila. Cell 163:1742–1755.

23. Dag U, Lei Z, Le JQ, Wong A, Bushey D, Keleman K (2019) Neuronal reactivation during post-learning sleep consolidates long-term memory in Drosophila. Elife 8.

24. Das S, Sadanandappa MK, Dervan A, Larkin A, Lee JA, Sudhakaran IP, Priya R, Heidari R, Holohan EE, Pimentel A, Gandhi A, Ito K, Sanyal S, Wang JW, Rodrigues V, Ramaswami M (2011) Plasticity of local GABAergic interneurons drives olfactory habituation. Proc Natl Acad Sci U S A 108:E646–654.

25. Davis RL (2011) Traces of Drosophila memory. Neuron 70:8–19.

26. de Belle JS, Heisenberg M (1994) Associative odor learning in Drosophila abolished by chemical ablation of mushroom bodies. Science 263:692–695.

27. Deisseroth K, Heist EK, Tsien RW (1998) Translocation of calmodulin to the nucleus supports CREB phosphorylation in hippocampal neurons. Nature 392:198–202.

28. Diekelmann S, Born J (2010) The memory function of sleep. Nat Rev Neurosci 11:114–126.

29. Dissel S, Melnattur K, Shaw PJ (2015) Sleep, Performance, and Memory in Flies. Curr Sleep Med Rep 1:47–54.

30. Donelson NC, Kim EZ, Slawson JB, Vecsey CG, Huber R, Griffith LC (2012) High-resolution positional tracking for long-term analysis of Drosophila sleep and locomotion using the “tracker” program. PLoS One 7:e37250.

31. Donlea JM (2019) Roles for sleep in memory: insights from the fly. Curr Opin Neurobiol 54:120–126.

32. Donlea JM, Ramanan N, Shaw PJ (2009) Use-dependent plasticity in clock neurons regulates sleep need in Drosophila. Science 324:105–108.

33. Donlea JM, Pimentel D, Miesenbock G (2014) Neuronal Machinery of Sleep Homeostasis in Drosophila. Neuron 81:1442.

34. Dudai Y (2012) The restless engram: consolidations never end. Annu Rev Neurosci 35:227–247.

35. Felsenberg J, Barnstedt O, Cognigni P, Lin S, Waddell S (2017) Re-evaluation of learned information in Drosophila. Nature 544:240–244.

36. Felsenberg J, Jacob PF, Walker T, Barnstedt O, Edmondson-Stait AJ, Pleijzier MW, Otto N, Schlegel P, Sharifi N, Perisse E, Smith CS, Lauritzen JS, Costa M, Jefferis G, Bock DD, Waddell S (2018) Integration of Parallel Opposing Memories Underlies Memory Extinction. Cell 175:709–722 e715.

37. Ferdenzi C, Poncelet J, Rouby C, Bensafi M (2014) Repeated exposure to odors induces affective habituation of perception and sniffing. Front Behav Neurosci 8:119.

38. Flyer-Adams JG, Rivera-Rodriguez EJ, Yu J, Mardovin JD, Reed ML, Griffith LC (2020) Regulation of Olfactory Associative Memory by the Circadian Clock Output Signal Pigment-Dispersing Factor (PDF). J Neurosci 40:9066–9077.

39. Ganguly-Fitzgerald I, Donlea J, Shaw PJ (2006) Waking experience affects sleep need in Drosophila. Science 313:1775–1781.

40. Gervasi N, Tchenio P, Preat T (2010) PKA dynamics in a Drosophila learning center: coincidence detection by rutabaga adenylyl cyclase and spatial regulation by dunce phosphodiesterase. Neuron 65:516–529.

41. Giese KP, Mizuno K (2013) The roles of protein kinases in learning and memory. Learn Mem 20:540–552.

42. Goel N, Rao H, Durmer JS, Dinges DF (2009) Neurocognitive consequences of sleep deprivation. Semin Neurol 29:320–339.

43. Griffith LC, Verselis LM, Aitken KM, Kyriacou CP, Danho W, Greenspan RJ (1993) Inhibition of calcium/calmodulin-dependent protein kinase in Drosophila disrupts behavioral plasticity. Neuron 10:501–509.

44. Handler A, Graham TGW, Cohn R, Morantte I, Siliciano AF, Zeng J, Li Y, Ruta V (2019) Distinct Dopamine Receptor Pathways Underlie the Temporal Sensitivity of Associative Learning. Cell 178:60–75 e19.

45. Hendricks JC, Finn SM, Panckeri KA, Chavkin J, Williams JA, Sehgal A, Pack AI (2000) Rest in Drosophila is a sleep-like state. Neuron 25:129–138.

46. Hige T, Aso Y, Modi MN, Rubin GM, Turner GC (2015) Heterosynaptic Plasticity Underlies Aversive Olfactory Learning in Drosophila. Neuron 88:985–998.

47. Honegger KS, Campbell RA, Turner GC (2011) Cellular-resolution population imaging reveals robust sparse coding in the Drosophila mushroom body. J Neurosci 31:11772–11785.

48. Inada K, Tsuchimoto Y, Kazama H (2017) Origins of Cell-Type-Specific Olfactory Processing in the Drosophila Mushroom Body Circuit. Neuron 95:357–367 e354.

49. Ito I, Ong RC, Raman B, Stopfer M (2008) Sparse odor representation and olfactory learning. Nat Neurosci 11:1177–1184.

50. Kelleher RJ, 3rd, Govindarajan A, Tonegawa S (2004) Translational regulatory mechanisms in persistent forms of synaptic plasticity. Neuron 44:59–73.

51. Kim YC, Lee HG, Han KA (2007) D1 dopamine receptor dDA1 is required in the mushroom body neurons for aversive and appetitive learning in Drosophila. J Neurosci 27:7640–7647.

52. Koh YH, Popova E, Thomas U, Griffith LC, Budnik V (1999) Regulation of DLG localization at synapses by CaMKII-dependent phosphorylation. Cell 98:353–363.

53. Krashes MJ, Keene AC, Leung B, Armstrong JD, Waddell S (2007) Sequential use of mushroom body neuron subsets during drosophila odor memory processing. Neuron 53:103–115.

54. Kuklin EA, Alkins S, Bakthavachalu B, Genco MC, Sudhakaran I, Raghavan KV, Ramaswami M, Griffith LC (2017) The Long 3’UTR mRNA of CaMKII Is Essential for Translation-Dependent Plasticity of Spontaneous Release in Drosophila melanogaster. J Neurosci 37:10554–10566.

55. Lei Z, Chen K, Li H, Liu H, Guo A (2013) The GABA system regulates the sparse coding of odors in the mushroom bodies of Drosophila. Biochem Biophys Res Commun 436:35–40.

56. Levin LR, Han PL, Hwang PM, Feinstein PG, Davis RL, Reed RR (1992) The Drosophila learning and memory gene rutabaga encodes a Ca2+/Calmodulin-responsive adenylyl cyclase. Cell 68:479–489.

57. Li F et al. (2020) The connectome of the adult Drosophila mushroom body provides insights into function. Elife 9.

58. Li X, Yu F, Guo A (2009) Sleep deprivation specifically impairs short-term olfactory memory in Drosophila. Sleep 32:1417–1424.

59. Lin AC, Bygrave AM, de Calignon A, Lee T, Miesenbock G (2014) Sparse, decorrelated odor coding in the mushroom body enhances learned odor discrimination. Nat Neurosci 17:559–568.

60. Livingstone MS, Sziber PP, Quinn WG (1984) Loss of calcium/calmodulin responsiveness in adenylate cyclase of rutabaga, a Drosophila learning mutant. Cell 37:205–215.

61. Lyons LC, Roman G (2009) Circadian modulation of short-term memory in Drosophila. Learn Mem 16:19–27.

62. Malik BR, Gillespie JM, Hodge JJ (2013) CASK and CaMKII function in the mushroom body alpha’/beta’ neurons during Drosophila memory formation. Front Neural Circuits 7:52.

63. Mao Z, Davis RL (2009) Eight different types of dopaminergic neurons innervate the Drosophila mushroom body neuropil: anatomical and physiological heterogeneity. Front Neural Circuits 3:5.

64. Mao Z, Roman G, Zong L, Davis RL (2004) Pharmacogenetic rescue in time and space of the rutabaga memory impairment by using Gene-Switch. Proc Natl Acad Sci U S A 101:198–203.

65. Marin EC, Jefferis GS, Komiyama T, Zhu H, Luo L (2002) Representation of the glomerular olfactory map in the Drosophila brain. Cell 109:243–255.

66. Masek P, Worden K, Aso Y, Rubin GM, Keene AC (2015) A dopamine-modulated neural circuit regulating aversive taste memory in Drosophila. Curr Biol 25:1535–1541.

67. McClelland JL, McNaughton BL, O’Reilly RC (1995) Why there are complementary learning systems in the hippocampus and neocortex: insights from the successes and failures of connectionist models of learning and memory. Psychol Rev 102:419–457.

68. Miller S, Yasuda M, Coats JK, Jones Y, Martone ME, Mayford M (2002) Disruption of dendritic translation of CaMKIIalpha impairs stabilization of synaptic plasticity and memory consolidation. Neuron 36:507–519.

69. Mitchell J, Smith CS, Titlow J, Otto N, van Velde P, Booth M, Davis I, Waddell S (2021) Selective dendritic localization of mRNA in Drosophila mushroom body output neurons. Elife 10.

70. Murthy M, Fiete I, Laurent G (2008) Testing odor response stereotypy in the Drosophila mushroom body. Neuron 59:1009–1023.

71. Olsen SR, Wilson RI (2008) Cracking neural circuits in a tiny brain: new approaches for understanding the neural circuitry of Drosophila. Trends Neurosci 31:512–520.

72. Owald D, Waddell S (2015) Olfactory learning skews mushroom body output pathways to steer behavioral choice in Drosophila. Curr Opin Neurobiol 35:178–184.

73. Owald D, Felsenberg J, Talbot CB, Das G, Perisse E, Huetteroth W, Waddell S (2015) Activity of defined mushroom body output neurons underlies learned olfactory behavior in Drosophila. Neuron 86:417–427.

74. Pech U, Pooryasin A, Birman S, Fiala A (2013) Localization of the contacts between Kenyon cells and aminergic neurons in the Drosophila melanogaster brain using SplitGFP reconstitution. J Comp Neurol 521:3992–4026.

75. Pellegrino R, Sinding C, de Wijk RA, Hummel T (2017) Habituation and adaptation to odors in humans. Physiol Behav 177:13–19.

76. Perez-Orive J, Mazor O, Turner GC, Cassenaer S, Wilson RI, Laurent G (2002) Oscillations and sparsening of odor representations in the mushroom body. Science 297:359–365.

77. Riemensperger T, Voller T, Stock P, Buchner E, Fiala A (2005) Punishment prediction by dopaminergic neurons in Drosophila. Curr Biol 15:1953–1960.

78. Schroll C, Riemensperger T, Bucher D, Ehmer J, Voller T, Erbguth K, Gerber B, Hendel T, Nagel G, Buchner E, Fiala A (2006) Light-induced activation of distinct modulatory neurons triggers appetitive or aversive learning in Drosophila larvae. Curr Biol 16:1741–1747.

79. Schwaerzel M, Monastirioti M, Scholz H, Friggi-Grelin F, Birman S, Heisenberg M (2003) Dopamine and octopamine differentiate between aversive and appetitive olfactory memories in Drosophila. J Neurosci 23:10495–10502.

80. Seidner G, Robinson JE, Wu M, Worden K, Masek P, Roberts SW, Keene AC, Joiner WJ (2015) Identification of Neurons with a Privileged Role in Sleep Homeostasis in Drosophila melanogaster. Curr Biol 25:2928–2938.

81. Sejourne J, Placais PY, Aso Y, Siwanowicz I, Trannoy S, Thoma V, Tedjakumala SR, Rubin GM, Tchenio P, Ito K, Isabel G, Tanimoto H, Preat T (2011) Mushroom body efferent neurons responsible for aversive olfactory memory retrieval in Drosophila. Nat Neurosci 14:903–910.

82. Semelidou O, Acevedo SF, Skoulakis EM (2018) Temporally specific engagement of distinct neuronal circuits regulating olfactory habituation in Drosophila. Elife 7.

83. Seugnet L, Suzuki Y, Vine L, Gottschalk L, Shaw PJ (2008) D1 receptor activation in the mushroom bodies rescues sleep-loss-induced learning impairments in Drosophila. Curr Biol 18:1110–1117.

84. Seugnet L, Suzuki Y, Thimgan M, Donlea J, Gimbel SI, Gottschalk L, Duntley SP, Shaw PJ (2009) Identifying sleep regulatory genes using a Drosophila model of insomnia. J Neurosci 29:7148–7157.

85. Shaw PJ, Cirelli C, Greenspan RJ, Tononi G (2000) Correlates of sleep and waking in Drosophila melanogaster. Science 287:1834–1837.

86. Simpson JH (2009) Mapping and manipulating neural circuits in the fly brain. Adv Genet 65:79–143.

87. Stewart BA, Atwood HL, Renger JJ, Wang J, Wu CF (1994) Improved stability of Drosophila larval neuromuscular preparations in haemolymph-like physiological solutions. J Comp Physiol A 175:179–191.

88. Sun F, Zhou J, Dai B, Qian T, Zeng J, Li X, Zhuo Y, Zhang Y, Wang Y, Qian C, Tan K, Feng J, Dong H, Lin D, Cui G, Li Y (2020) Next-generation GRAB sensors for monitoring dopaminergic activity in vivo. Nature methods 17:1156–1166.

89. Suzuki-Sawano E, Ueno K, Naganos S, Sawano Y, Horiuchi J, Saitoe M (2017) A Drosophila ex vivo model of olfactory appetitive learning. Sci Rep 7:17725.

90. Tanaka NK, Tanimoto H, Ito K (2008) Neuronal assemblies of the Drosophila mushroom body. J Comp Neurol 508:711–755.

91. Tomchik SM, Davis RL (2009) Dynamics of learning-related cAMP signaling and stimulus integration in the Drosophila olfactory pathway. Neuron 64:510–521.

92. Tully T, Quinn WG (1985) Classical conditioning and retention in normal and mutant Drosophila melanogaster. J Comp Physiol A 157:263–277.

93. Turner GC, Bazhenov M, Laurent G (2008) Olfactory representations by Drosophila mushroom body neurons. J Neurophysiol 99:734–746.

94. Ueno K, Naganos S, Hirano Y, Horiuchi J, Saitoe M (2013) Long-term enhancement of synaptic transmission between antennal lobe and mushroom body in cultured Drosophila brain. J Physiol 591:287–302.

95. Ueno K, Suzuki E, Naganos S, Ofusa K, Horiuchi J, Saitoe M (2017) Coincident postsynaptic activity gates presynaptic dopamine release to induce plasticity in Drosophila mushroom bodies. Elife 6.

96. Ueno K, Morstein J, Ofusa K, Naganos S, Suzuki-Sawano E, Minegishi S, Rezgui SP, Kitagishi H, Michel BW, Chang CJ, Horiuchi J, Saitoe M (2020) Carbon Monoxide, a Retrograde Messenger Generated in Postsynaptic Mushroom Body Neurons, Evokes Noncanonical Dopamine Release. J Neurosci 40:3533–3548.

97. Wang Y, Mamiya A, Chiang AS, Zhong Y (2008) Imaging of an early memory trace in the Drosophila mushroom body. J Neurosci 28:4368–4376.

98. Wiggin TD, Goodwin PR, Donelson NC, Liu C, Trinh K, Sanyal S, Griffith LC (2020) Covert sleep-related biological processes are revealed by probabilistic analysis in Drosophila. Proc Natl Acad Sci U S A 117:10024–10034.

99. Wilson DA (1998) Habituation of odor responses in the rat anterior piriform cortex. J Neurophysiol 79:1425–1440.

100. Wilson DA (2009) Olfaction as a model system for the neurobiology of mammalian short-term habituation. Neurobiol Learn Mem 92:199–205.

101. Wilson DA, Linster C (2008) Neurobiology of a simple memory. J Neurophysiol 100:2–7.

102. Wong AM, Wang JW, Axel R (2002) Spatial representation of the glomerular map in the Drosophila protocerebrum. Cell 109:229–241.

103. Yamagata N, Hiroi M, Kondo S, Abe A, Tanimoto H (2016) Suppression of Dopamine Neurons Mediates Reward. PLoS Biol 14:e1002586.

104. Yao Z, Macara AM, Lelito KR, Minosyan TY, Shafer OT (2012) Analysis of functional neuronal connectivity in the Drosophila brain. J Neurophysiol 108:684–696.

105. Yu D, Akalal DB, Davis RL (2006) Drosophila alpha/beta mushroom body neurons form a branch-specific, long-term cellular memory trace after spaced olfactory conditioning. Neuron 52:845–855.

106. Zhang X, Noyes NC, Zeng J, Li Y, Davis RL (2019) Aversive Training Induces Both Presynaptic and Postsynaptic Suppression in Drosophila. J Neurosci 39:9164–9172.

107. Zhao X, Lenek D, Dag U, Dickson BJ, Keleman K (2018) Persistent activity in a recurrent circuit underlies courtship memory in Drosophila. Elife 7.

